# Penetratin inhibits α-synuclein fibrillation and improves locomotor functions in mice model of Parkinson’s disease

**DOI:** 10.1101/2022.06.24.497475

**Authors:** Arpit Gupta, Priyanka Singh, Arpit Mehrotra, Ankur Gautam, K. Srividya, Rajlaxmi Panigrahi, Shubham Vashishtha, Jasdeep Singh, Gagandeep Jaiswal, Krishna Upadhayay, Signe Andrea Frank, Janni Nielsen, Samir Kumar Nath, Neeraj Khatri, Daniel E. Otzen, G.P.S. Raghava, Anil Koul, Bishwajit Kundu, Ashutosh Kumar, Aamir Nazir, Deepak Sharma

## Abstract

Parkinson’s disease (PD) is the second most common neurodegenerative disease. The presence of lewy bodies, primarily consisting of α-synuclein (α-syn) aggregates is one of the common features seen in the substantia nigra region of the brain in PD patients. The disease remains incurable and only symptomatic relief is available. We screened various cell-penetrating peptides and reveal that penetratin is a potent inhibitor of α-syn aggregation *in-vitro*, and significantly improved locomotor coordination in mice models of PD *in-vivo*. The peptide inhibits α-syn aggregation in vitro as well as in yeast, and *C.elegans* models. We further made a cyclic derivative of penetratin by disulfide coupling of N- and C-terminal cysteine residues. Both penetratin and its cyclized derivative interact with α-syn. NMR studies show that both linear as well as cyclic derivative interact at the acidic C-terminal tail of the protein. Similar to penetratin, its cyclic derivative inhibited α-syn aggregation in the *C.elegans* model of Parkinson’s disease, and also improved worm motility. Molecular Dynamics studies show that penetratin interacts with α-synuclein and prevents its conformational transition from disordered into β-sheet rich structure. The therapeutic efficacy of penetratin was further confirmed in a transgenic mice model of the disease, wherein penetratin treatment over a period of 90 days improved locomotor coordination, and halted disease progression. Overall, the present work provides a potent therapeutic agent that could be further explored in the management of PD.

## Introduction

Parkinson’s disease (PD), the second most common neurodegenerative disease is characterized by the neuronal accumulation of Lewy bodies in the substantia nigra region of the brain (Mahul-Mellier et al., 2020). The Lewy bodies are protein-rich intracellular inclusions primarily composed of fibrillar α-syn aggregates. The cytoplasmic accumulation of these aggregates in dopaminergic neurons is associated with neuronal toxicity which results in deficiency of dopamine required for motor movements and reward responses in the brain. Thus, Parkinson’s patients show typical symptoms of movement disorders such as bradykinesia, resting tremor, and also posture instability at later stages of the disease. In addition to PD, α-syn aggregation is also involved in various other human diseases collectively known as α-synucleinopathies such as Dementia with Lewy Bodies (DLB) and Multiple System Atrophy (MSA). Though symptomatic relief is available, finding an effective cure against α-synucleinopathies still remains a substantial challenge.

α-syn is a member of the synuclein family of proteins abundantly present in presynaptic terminals of neuronal tissues. The synucleins are characterized by the presence of 5 or 6 imperfect repeats of motif KTKEGV followed by a central hydrophobic non-amyloid β component (NAC) region and an acidic carboxyl-terminal tail. Mutations (such as A30P, A53T and E46K) in SNCA, the gene that encodes (α- syn, are associated with familial forms of PD (Dawson and Dawson, 2003; Krüger et al., 1998; Polymeropoulos et al., 1997; Zarranz et al., 2004). Multiple functions have been associated with α-syn such as regulation of neuronal plasticity, polymerization of purified tubulin into microtubules and formation of SNARE complexes (protein complexes critical for release of neurotransmitters) (Alim et al., 2004; Chandra et al., 2005; Jin and Clayton, 1997). α-syn is considered to be non-essential for cellular survival as mice that lack α-syn exhibit normal behaviour and possess normal dopaminergic bodies though a reduction in striatal dopamine has been observed (Abeliovich et al., 2000). Cellular toxicity is believed to be caused by protofibrils that are formed during the conformational transition of native α-syn into fibrils (Bucciantini et al., 2002; Cascella et al., 2021), and thus strategies to block the process of fibrillation is one of the extensively targeted approaches in search of therapeutics against PD.

Current approaches employed in reducing α-syn associated toxicity include designing small-molecule based inhibitor/s of fibrillation, reduced expression or enhanced degradation of the protein (Di Giovanni et al., 2010; Lashuel et al., 2013). As α-syn is a disordered protein, the rational design of its binders and inhibitors of aggregation has been a relatively difficult task. Thus, the identification of suitable inhibitors of α-syn fibrillation has primarily been carried out by screening large libraries of small molecules. Attempts have been made to further enhance the efficacy of inhibitors by increasing their ability to cross the blood-brain barrier using approaches based on nanoparticles or cell-penetrating peptides (CPPs) (Zhang et al., 2021).

CPPs have been extensively explored for their ability to deliver bimolecular cargoes inside the cells. The CPPs translocate across cellular membranes without damaging or having any significant adverse effect on cellular membranes. Based on their physicochemical properties, CPPs are broadly classified into three classes (i) cationic (ii) amphipathic and (iii) hydrophobic (Kang et al., 2019; Skwarczynski and Toth, 2019). Though the mechanism of cell entry remains under debate, both passive permeation across cell membranes and endocytosis are widely believed to be the underlying basis of CPPs translocation (Feni and Neundorf, 2017). Many of the CPPs are able to cross the blood-brain barrier (BBB), and thus have been used as cargo to transport conjugated desired small-molecule or proteins to the brain (Qin et al., 2011; Qin et al., 2012). Considering their ability to cross BBB, such peptides could be of potential use in the treatment of brain-related disorders.

Though various compounds have been shown to inhibit α-syn fibrillation *in vitro*, their ability to cross the BBB or to penetrate through the cell membrane to access intracellular aggregates remains a significant challenge. In the present study, we screened some of the well-studied CPPs for their ability to inhibit α-syn fibrillation. Among several CPPs that were screened, we show that penetratin efficiently inhibited α-syn fibrillation at sub-stoichiometric concentration. Penetratin binds to monomeric α-syn with a K_D_ of 245 nM. The molecular dynamics simulation studies confirmed that penetratin interacts at the C-terminal of α-syn and subsequently inhibits its conformational transition to β-sheet formation. The interaction was also confirmed by NMR analysis. Furthermore, penetratin treatment showed reduced α- syn aggregation in the *C.elegans* model of PD and ameliorated motor dysfunctions in the transgenic mice model of PD.

## Results

### Penetratin inhibits α-syn aggregation *in vitro*

The cytoplasmic accumulation of α-syn amyloid aggregates is one of the common features observed in PD. Though several inhibitors of α-syn fibrillation have been identified, the inability of many of such molecules to cross the cellular membrane has been a major limitation in their use as an effective therapy. Thus, we screened various previously known extensively used CPPs for their ability to inhibit α-syn fibrillation. Table 1 describes the basis of the selection and sequence of the selected peptides. The peptides were primarily selected on the basis of predicted relatively higher blood brain barrier score, lower haemolytic activity and toxicity (Table 1). Among amphipathic peptides, we selected well-characterized and widely used pVEC, ARF, MAP and TP-10 while TAT, R8, IMT-P8, SynB3 and Penetratin were selected from the cationic class (Stalmans et al., 2015; Zhang et al., 2021). Peptides from the hydrophobic class were not selected due to their limited solubility and high propensity to aggregate.

**Table 1:**
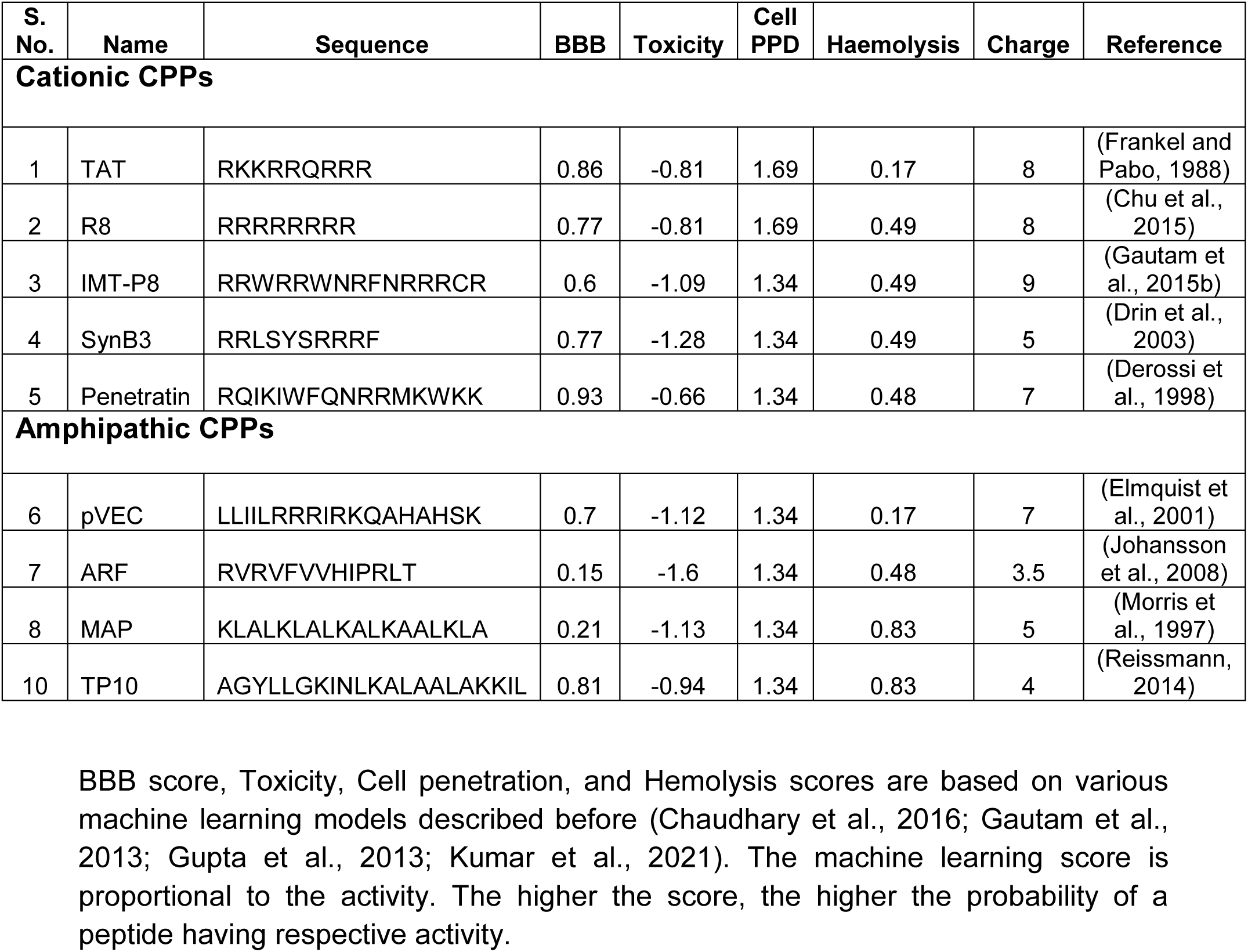
Cell Penetrating Peptides (CPPs) screened in the present study with predicted BBB, toxicity, cell penetrating ability and haemolytic activity score.

TAT and penetratin are among the first few peptides shown to have cell penetrating ability(Derossi et al., 1998; Frankel and Pabo, 1988; Vivès et al., 1997). TAT is derived from Transcription transactivating protein (TAT) encoded by human immunodeficiency virus type 1 (HIV-1), and penetratin from Antennapedia homeodomain (Frankel and Pabo, 1988; Prochiantz, 1999). Both TAT and penetratin have been extensively used for the intracellular delivery of a variety of cargoes both in vitro and in vivo settings (Milletti, 2012; Shin et al., 2014). SynB3 is cationic peptide capable of crossing the cellular membranes without compromising its integrity(Drin et al., 2003). pVEC is an 18 amino acid peptide derived from murine vascular endothelial-cadherin protein (Elmquist et al., 2001). TP-10 and R8 are other extensively used CPPs (Soomets et al., 2000; Wender et al., 2000). IMT-P8 is an arginine-rich natural protein-derived peptide and has been shown to deliver cargoes in cell lines as well as into the skin (Gautam et al., 2016; Gautam et al., 2015b). MAP is 18-mer artificially designed peptide with both hydrophobic and hydrophilic residues which enables its cellular internalization with a non-endocytic pathway (Scheller et al., 2000). ARF is derived from the N-terminal part of tumor suppressor protein p14ARF (Johansson et al., 2008).

The peptides were examined for their ability to inhibit α-syn fibrillation using a well-established Thioflavin T (ThT) based fluorescence assay (Figure 1A). ThT binds specifically to fibrillar aggregates which leads to an increase in its fluorescence at 482nm. 400 µM of α-syn, in the presence and absence of the desired peptide,was incubated at 37°C, and fibrillation was monitored at regular time intervals. In the absence of any peptide, the α-syn fibrillation showed a lag phase of about ∼2h followed by a gradual increase in ThT fluorescence which levelled off around 6h. The addition of different peptides had different effect on α-syn fibrillation. TP-10, TAT, ARF, and SynB3 increased the lag phase of fibrillation whereas IMT-P8, pVEC, R8 and MAP reduced the lag phase (Table 2).The CPP IMT-P8 not only reduced the lag phase but also a concomitant increase in the rate of elongation and amplitude at saturation phase. Remarkably, the addition of an equimolar amount of penetratin completely suppressed any increase in ThT fluorescence during the time course (8h) of the fibrillation reaction. Overall among the CPPs tested, only penetratin showed strong inhibitory effect on α-syn fibrillation.

**Figure 1:**
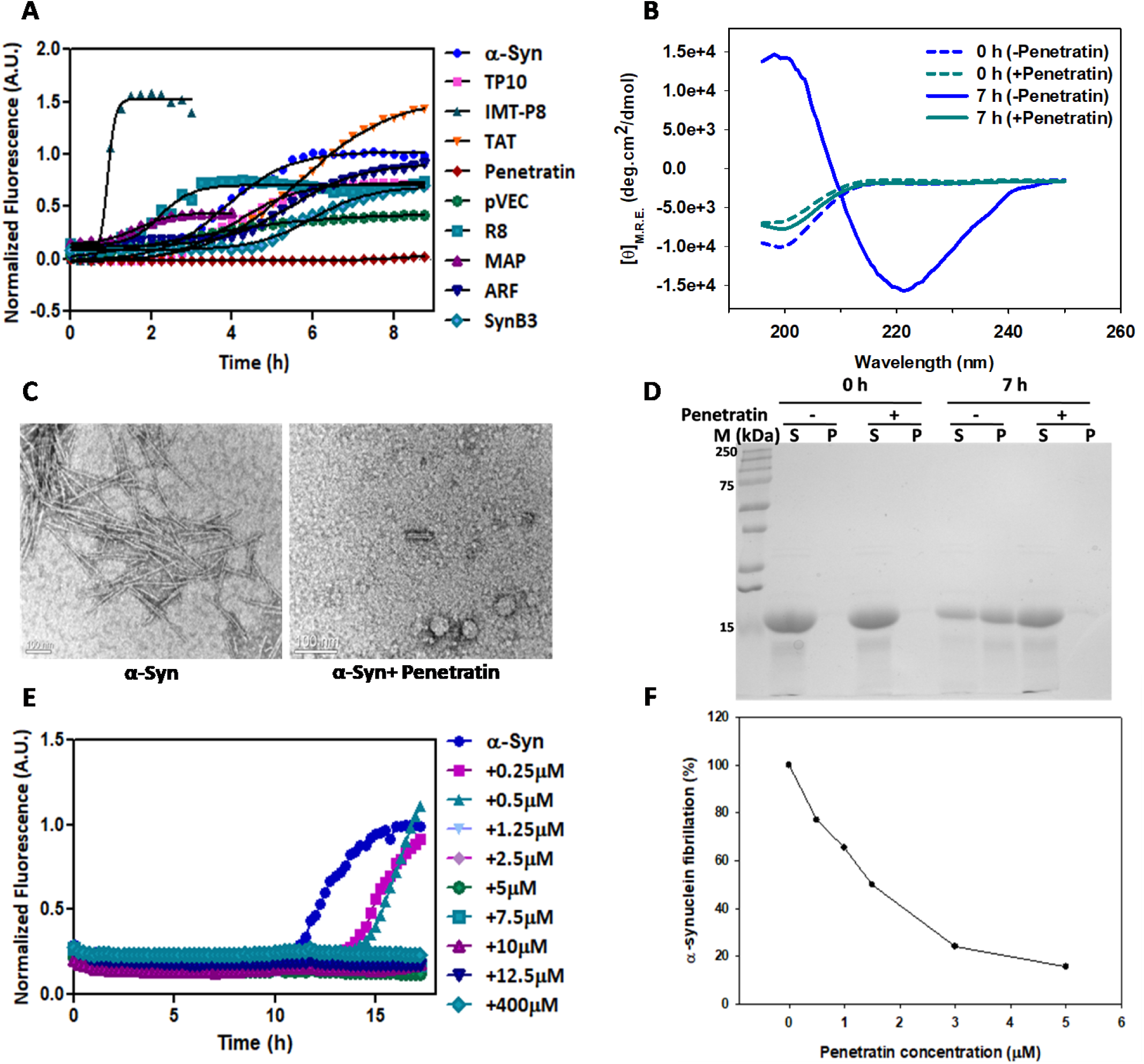
Penetratin inhibits α-Syn fibrillation. **(A)** α-Syn (400µM), in the presence or absence of equimolar concentrations of indicated peptides, was incubated at 37°C. The fibrillation was monitored by increase inThTfluorescenceintensity.**(B)**The CD study was carried out with samples obtained from fibrillation reaction at 0h and 7h of incubation at 37°C. **(C)**TEM images of samples obtained as in panel B.**(D)**The fibrillation reaction mixture was fractionated. The supernatant and pellet were loaded onto 15% SDS-PAGE. **(E)**The α- synfibrillation was carried out with varying concentrations of penetratin at 37°C for 18h and**(F)**fractionated to separate the pellet fraction. The pellet fraction loaded onto 15% SDS-PAGE, and band intensity for α-synwas measured using UN-SCANIT. Shown is the normalized band intensityof α-Syn on SDS-PAGE versus penetratin concentration.

**Table 2:** Calculated kinetic parameters for screened CPPs against α-Syn fibrillation

| Serial no. | Peptides | $T_{1/2}$ (h) | $T_0$ (Lag time) (h) |
| --- | --- | --- | --- |
| 1. | α-Syn | 3.83 | 2.26 |
| 2. | TP10 | 4.05 | 2.45 |
| 3. | IMT-P8 | 0.92 | 0.72 |
| 4. | TAT | 5.68 | 3.4 |
| 5. | Penetratin | 8.13 | 7.27 |
| 6. | pVEC | 2.68 | 0.7 |
| 7. | R8 | 2.21 | 1.42 |
| 8. | MAP | 1.62 | 0.78 |
| 9. | ARF | 5.46 | 3.74 |
| 10. | SynB3 | 5.97 | 4.55 |

During fibrillation, α-syn is known to undergo a transition from an intrinsically disordered to β-sheet-rich state. Thus, we examined the effect of penetratin on conformational changes in α-syn. The α-syn fibrillation was carried out until the plateau (7h) was attained and CD spectra were recorded before and after removing the aggregates. As shown in Figure 1B, at 0h in the presence and absence of the peptide, α-syn showed CD spectra characteristic of an intrinsically disordered protein with a minimum around 200 nm and little signal above 210 nm. To obtain CD spectra of α-syn fibrils, the samples after 7h of fibrillation (early saturation phase) were collected and spun at 4000g for 1 min to remove visible aggregates. The CD spectra monitored after removal of aggregates was subtracted from that obtained with total reaction mixture to finally attain CD spectra for aggregates alone. As expected, the α-syn fibrils alone showed CD spectra characteristic of a β-sheet rich protein. Interestingly, no visible aggregates were observed even after 7h of fibrillation reaction in the presence of penetratin, and CD spectra observed was similar to as that observed at 0h, thus suggesting that penetratin inhibited the conformational transition of α-syn from a disordered state to β-sheet rich structural formation.

We further examined the effect of penetratin on α-syn fibril formation using TEM. The α-syn fibrillation reaction was performed as described previously. The reaction mixture was then placed onto copper grids followed by negative staining with phosphotungstic acid and observed under the electron microscope. As depicted in Figure 1C, only α-syn alone formed fibrils, and no fibrils were observed when penetratin was added to the reaction mixture.

Furthermore, the extent of α-syn aggregation in the presence and absence of penetratin was evaluated by SDS-PAGE. Following 7h of α-syn incubation at 37°C, with and without penetratin, the reaction was fractionated into supernatant and pellet containing aggregates, and fractions were boiled with 2% SDS prior to SDS-PAGE analysis (Figure 1D). Prior to initiation of fibrillation (at 0h), α-syn was primarily detected in supernatant fraction regardless of the presence of penetratin. However, post 7h, a significant amount (∼50%) of α-syn was observed in the pellet fraction in the absence of penetratin. Interestingly, α-syn pre-incubated with penetratin during the fibrillation process was found to be primarily present in soluble fraction even after 7h of incubation at 37°C, further confirming that the peptide inhibits α-syn aggregates formation.

We next evaluated the optimum concentration/s of penetratin in significantly inhibiting α-syn aggregates formation. Therefore, fibrillation reactions were performed in the absence and presence of varying concentrations of penetratin. Penetratin was added at increasing concentrations ranging from 0.25μM to 400μM into the reaction mixture containing α-syn (400µM), and fibrillation was monitored over a period of time (∼18h) in the presence of ThT dye (Figure 1E). The ThT assay revealed that although α-syn formed amyloid aggregates at lower concentrations of the penetratin (0.25μM and 0.5μM), the lag phase preceding fibrillation increased from ∼11h to 13h. Further, an increase in penetratin concentration to 1.25μM or more completely inhibited α-syn fibrillation as evidenced by lack of any increase in ThT fluorescence.

This concentration dependent effect of penetratin was confirmed by SDS-PAGE analysis. As such, varying concentrations (ranging from 0.5μM to 5μM) of penetratin were incubated with α-syn at 37°C. After 7 h, the reaction mixture was fractionated into supernatant and pellet fractions. The individual pellet fractions containing insoluble aggregates were boiled with 2% SDS and then loaded onto SDS-PAGE. Figure 1F depicts relative quantification of the protein band intensity as determined from SDS-PAGE analysis. As expected, at 0.5μM of penetratin most of α-syn was observed in pellet fraction. Further, increase in penetratin concentration to 1.5μM, 3μM and 5μM lead to decrease of α-syn in pellet fraction by 50%, 20% and 12% respectively, suggesting that an increase in penetratin led to a corresponding decrease in the α-syn aggregation. (Figure S1).

### α-syn retains a random coil in the presence of penetratin

The above data shows that penetratin is a potent inhibitor of α-syn fibrillation. In order to understand the underlying mechanism of the inhibitory effect, we carried out molecular dynamic simulations. Firstly, we studied thermally induced transitions of α-syn to β-rich metastable states. We hypothesized that these initial transitions might act as trigger points for the aggregation of the full-length protein. Simulation in the presence of the penetratin were carried out to assess possible interference with these transitions. Consequently, free energy landscapes were constructed from simulations of α-syn alone and in the presence of penetratin (Figure 2). These were projected onto the percentage coil and β-content of the system for respective simulations. In simulations of α-syn alone, the lowest free energy basin corresponded to a conformation with ∼60% coil and ∼25% of β-content (Figure 2A). The corresponding α-syn structure also showed the presence of anti-parallel β- sheets (Figure 2B). On the contrary, the lowest free energy basin for α-syn in the presence of the penetratin showed relatively higher coil content (∼75%) and lower β- sheet content (∼5%) (Figure 2C). Extraction of the corresponding α-syn structure showed the absence of an organized anti-parallel sheet arrangement (Figure 2D). Overall, the molecular dynamic simulations showed that penetratin inhibits the conformational transition of α-syn from a disordered state to a β-sheet rich structure.

**Figure 2:**
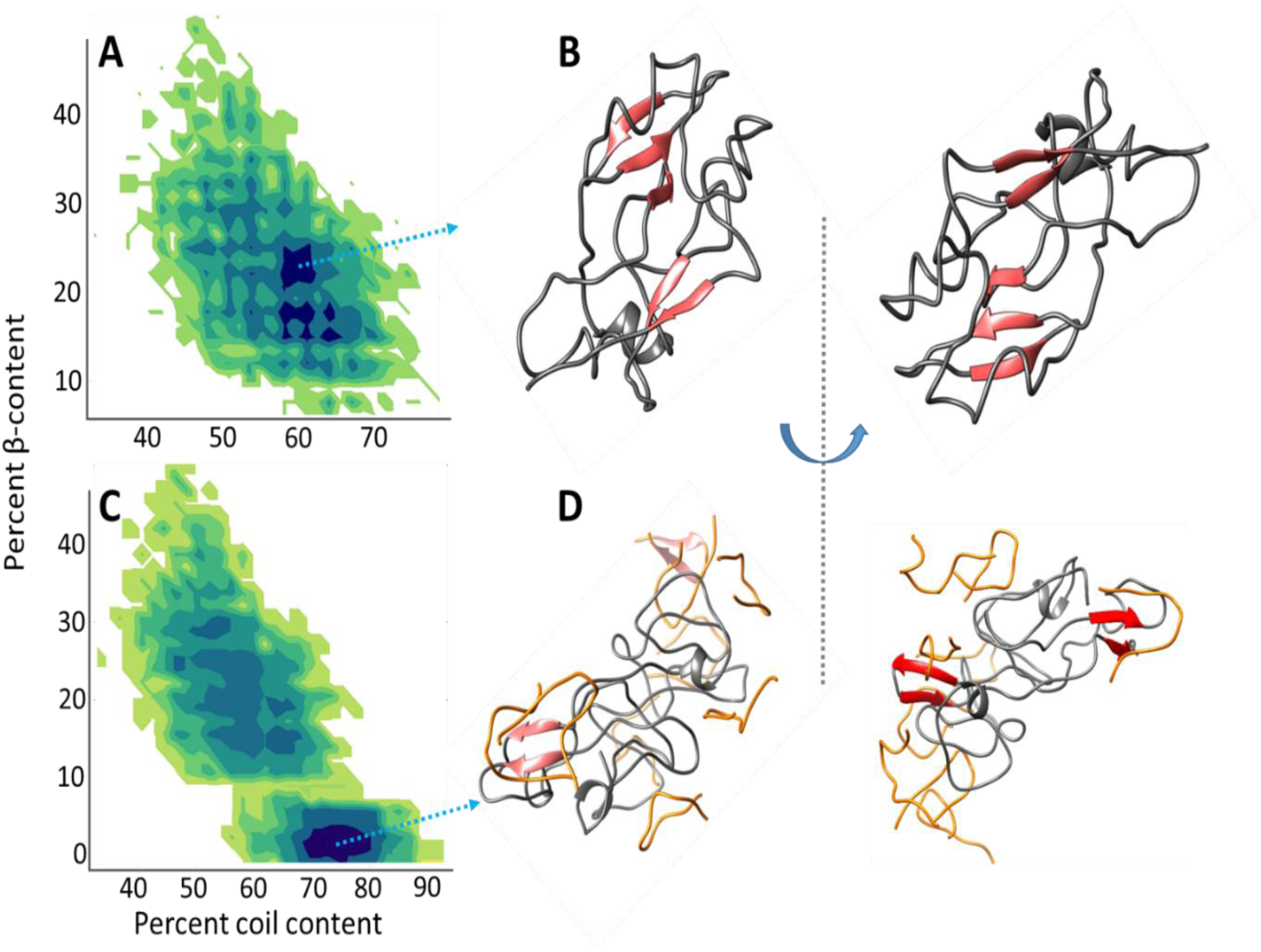
REMD simulations to understand α-syn conformational transition in the presence and absence of Penetratin. Free energy landscapes (kcal/mol) projected as a function of total coil content vs total β-content during replica exchange MD simulation and their corresponding structures extracted from lowest free energy basin frame in**A, B)**α-syn alone and **C, D)**α-syn with penetratin.

### Penetratin derivatives also inhibits α-syn fibrillation

It is known that cyclization enhances the peptide stability and resistance against proteolysis (Clark et al., 2005; Ngambenjawong et al., 2016; Ngo et al., 2020). In order to examine whether such modifications would have any adverse effect on inhibition of α-syn fibrillation, we designed a cyclic derivative of penetratin ‘cyclic-penentratin”. The cyclization of the peptide was achieved through addition of cys residues at the N- and C-termini of penetratin, followed by oxidation of the free cysteine residues to form an intramolecular disulfide bridge (cysteine).The designed peptide derivative was examined for its ability to inhibit α-syn fibrillation using ThT assay as described above (Figure 3A). As shown above, penetratin completely inhibited α-syn fibrillation. Similar to as with penetratin, there was no observed increase in ThT intensity in the presence of the cyclic-penetratin, indicating that it inhibited α-syn fibrillation as well. The reaction mix at 4h was further examined under TEM, wherein no α-syn fibrils were observed in reaction containing both α-syn and peptide derivative (Figure 3B).

**Figure 3:**
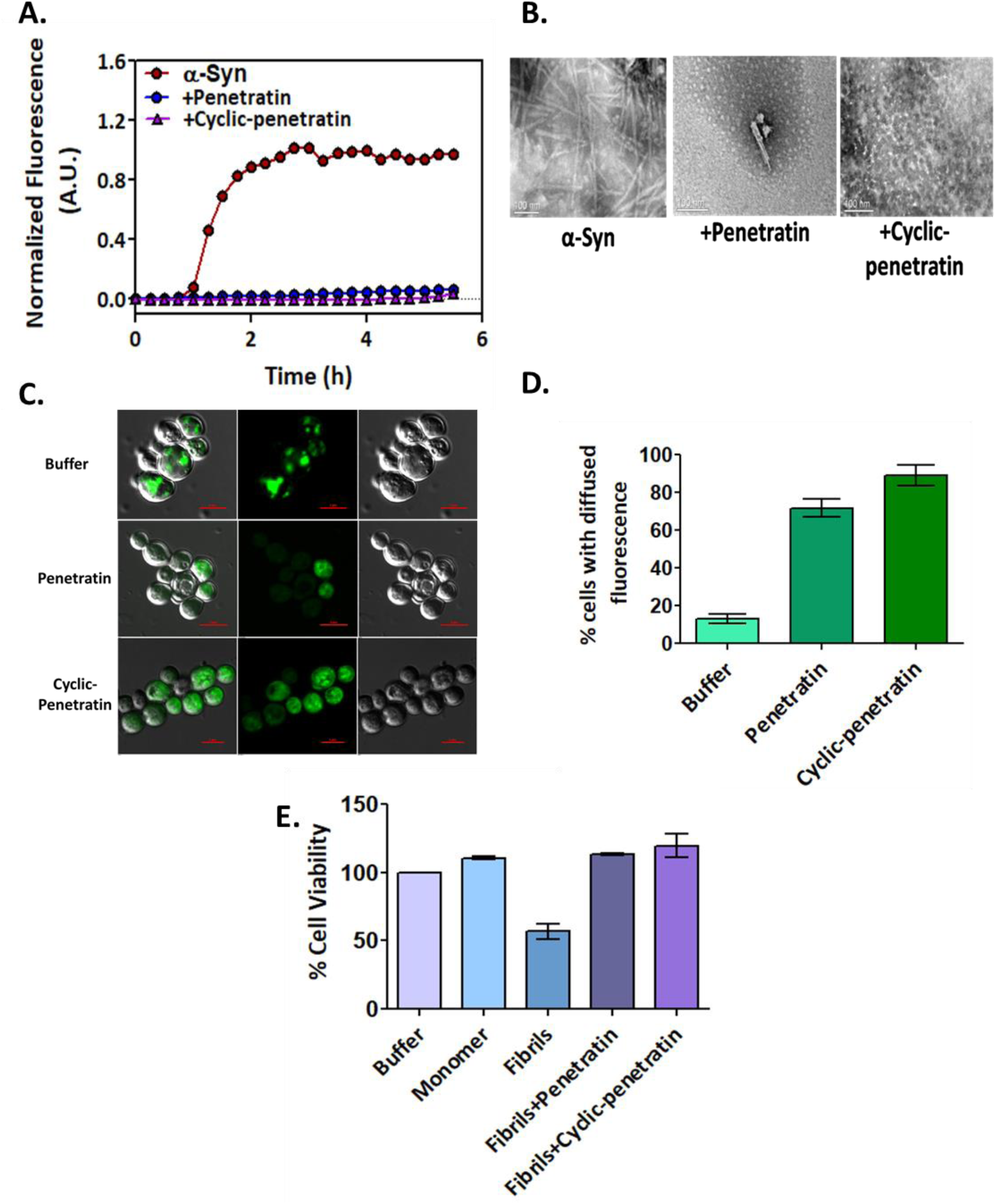
Both penetratin and its cyclic derivative inhibit α-syn fibrillation. **(A)** α-Syn (400µM) fibrillation was carried out in the presence or absence of equimolar concentrations of penetratin or cyclic-penetratin. The changes inThT fluorescence intensity (normalized) over time is shown. **(B)**TEM images of samples obtained at 4h of the fibrillation. **(C)** The *S. cerevisiae* strain (SY246) expressing α-Syn-GFP was incubated with penetratin or cyclic- penetratin for 60 min. The α-syn aggregates are seen as green punctae. Minimum of 100 cells expressing α-syn-GFP were examined, and**(D)**frequency of cells showing diffused α-syn- GFP fluorescence was quantified. (E) The α-syn fibrillation was carried out in the presence and absence of penetratin and cyc-penetratin. The samples were sonicated, and reaction mixture (10µM) incubated with SH-SY5Y for 24h at 37°C. The cellular viability was examined using MTT assay.

We further examined the ability of these derivatives to inhibit α-syn aggregation in a well-established yeast model of PD. Overexpression of α-syn from two genomically integrated SNCA-GFP (encoding α-syn-GFP) under galactose inducible promoter leads to aggregation of α-syn which are observed as green punctae under a fluorescence microscope. To examine whether designed peptide derivative could inhibit α-syn aggregation, we cultured yeast strain (SY246) in galactose inducible growth media for 12h. The cells were then incubated with peptide derivative (200µM) for 60 min and examined under confocal microscope. As seen in Figure 3C, 3D and S5, in the absence of the peptide derivative ∼90% of yeast cells showed α-syn aggregates. In contrast, most of the cells (>70%) co-incubated with either penetratin or cyclic-penetratin for 60 min showed diffused α- syn-GFP fluorescence, indicating inhibition of α-syn aggregation (Figure 3C and 3D). The above results were also investigated *in vitro* for validating the efficacy of penetratin and its derivative in reducing cellular toxicity in SHSY5Y neuroblastoma cells by MTT viability assay. The fibrillation reaction was carried out in the presence and absence of the peptide or its derivative until log phase was achieved, and further 10 µM of reaction mixture was incubated with SHSY5Y cells for 24h followed by MTT assay (Figure 3E).The treatment of cells with α-syn did not affect their viability. In comparison, cellular viability was reduced to ∼60% when incubated with α-syn fibrils obtained at log phase of fibrillation reaction. Interestingly, addition of penetratin or cyclic-penetratin in the fibrillation reaction restored cellular viability in SHSY5Y cells near to that observed in controls, thus strongly suggesting that the peptides reduced cellular stress against α-syn fibril associated toxicity.

### Penetratin and cyclic-penetratin interact with α-syn

We further investigated the ability of penetratin and cyclic-penetratin to interact with α-syn using microscale thermophoresis (MST). Peptides were serially diluted and incubated with a constant concentration of RED-NHS labelled α-syn for thermophoresis study. Figure 4 shows a dose-response curve obtained by plotting MST signal (ΔF_norm_) at varying peptide concentrations. We observed a decrease in normalized fluorescence intensity upon increase in either of the peptide concentrations, which indicates an increased diffusion of the bound complex as compared to free α-syn. The dose-response curves yielded a K_D_ of 245.4 ± 9.1 µM and 377.2 ± 21.0 µM for linear penetratin and cyclic-penetratin respectively, indicating relatively lower affinity of α-syn with the cyclic peptide.

**Figure 4:**
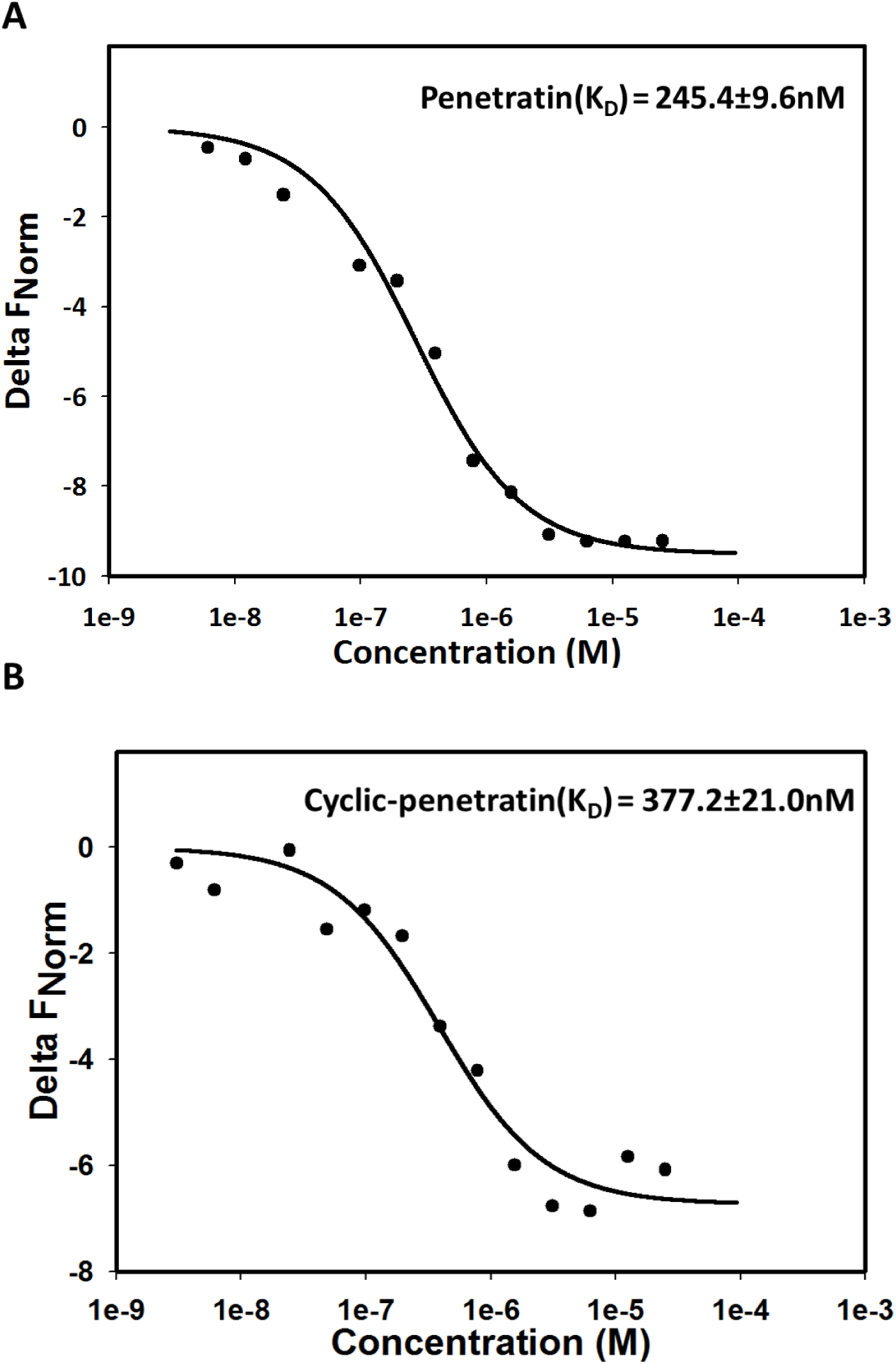
Penetratin and cyclic-penetratin interact with α-Syn. Penetratin and cyclic- penetratinat concentration ranging from 100µM to 3.05nM were incubated with constant amount of α-syn (40nM) and their binding was measuredusing microscale thermophoresis. Shown is the change in normalized fluorescence with peptide concentration.

### NMR studies show α-syn residues that interact with penetratin and its cyclized derivative

To investigate α-syn residues involved in interaction with the penetratin and cyclic-penetratin, ^1^H-^15^N Heteronuclear Single Quantum Coherence (HSQC) spectroscopy experiment was conducted for α-syn alone (Figure 5A) or in the presence of various molar equivalents of cyclic-penetratin (Figure 5B) and Penetratin (Figure 5C). The disordered nature of α-syn is not altered even in the presence of high concentrations of the peptides, as indicated by the narrow dispersion of the cross amide peaks in the spectra. However, the cross amide-peaks of the C-terminal residues (Q109-A140) of α-syn exhibit significant changes in the chemical shifts along with decrease in the intensity in the presence of both peptides (Figure 5F, G). This indicates that nature of the interaction is in the intermediate range of the NMR regime.The binding of peptides to the C-terminus of α-synuclein could be explained due to the presence of electrostatic interaction between the lysine and arginine-rich peptides and the negatively charged amino acid rich C-terminus of α-syn.

**Figure 5:**
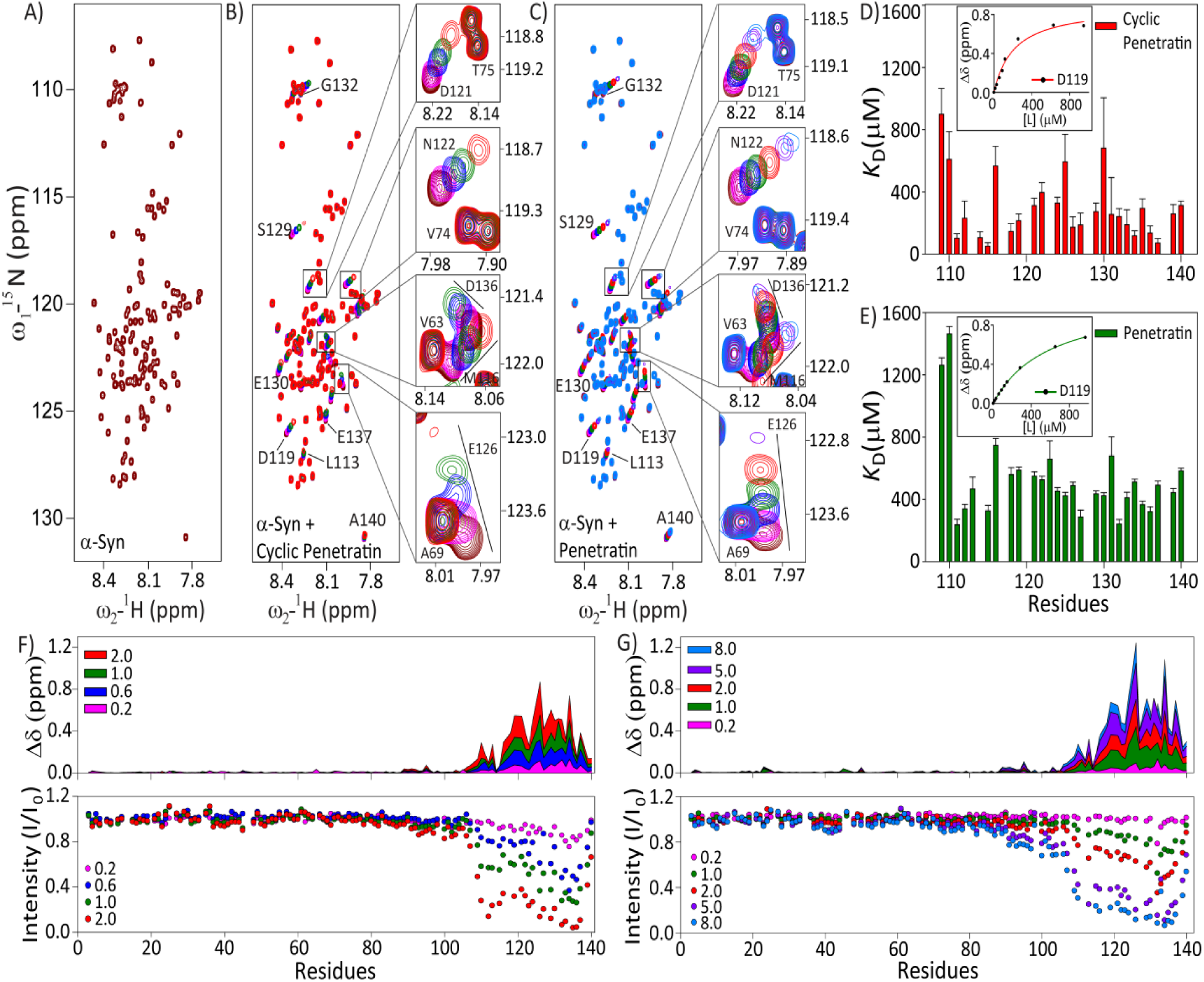
^1^H-^15^N Heteronuclear Single Quantum Coherence spectra of A) α-Syn. (maroon) and in presence of different concentrations (molar equivalents) of **B)** Cyclic- penetratin(magenta: 0.2; blue: 0.6; green: 1.0; and red: 2.0) and **C)**Penetratin(magenta: 0.2; blue: 0.6; green: 1.0; red: 2.0; mauve: 5.0; and sky-blue: 8.0). Residues exhibiting significant perturbations in the chemical shift have been highlighted in the insets and labeled in spectra. The direction of peak movement for cyclic-penetratin and penetratin is from maroon to red, and maroon to sky-blue respectively.Dissociation constant (*K*_D_) of the C-terminus residues of α-Syn exhibiting significant perturbations in chemical shifts in presence of the **D)** cyclic- penetratin and **E)**penetratinare calculated by the formula described in methodology. Chemical shift perturbations (Δδ) and Intensity (I/I_0_) profile of cross amide peaks obtained from ^1^H-^15^N HSQC spectra of α-Syn in presence of different molar equivalents of **F)** Cyclic-penetratin and **G)**Penetratin.

Although there is a perturbation in the conformation of α-syn at the C-terminal in the presence of either of the peptides, the strength of interaction varies. Figure 5F- G shows the chemical shift perturbation and intensity profile of cross amide peaks of α-syn in the presence of various molar equivalents of both the peptides. Cyclic- penetratin shows a great decrease in intensity at 2.0 molar equivalents along with chemical shift perturbation (CSP) in comparison to that of penetratin. The dissociation constant (*K*_D_) obtained for the C-terminal residues (Figure 5 D-E and Table S1) involved in the interaction of α-syn with these peptides based on chemical shift perturbation was found to be 297.6 ± 41.2 µM and 528.4 ± 52.6 µM for cyclic- penetratin and penetratin respectively (Figure S2 and Table S1).

### The C-terminal part of penetratin is particularly important for α-syn recognition

To investigate which region of penetratin are particularly important for interactions with α-syn, we prepared a peptide array displaying different variants of penetratin. The array was then exposed to fluorophore-labelled α-syn and the extent of binding to each peptide was quantified by scanning and normalized to the binding to wildtype (wt) penetratin. We focused primarily on two approaches: Ala-scanning, in which each of penetratin’s 16 residues were replaced by Ala, one residue at a time, and terminal truncation in which residues were removed in a cumulative fashion from either the C-terminus or the N-terminus. The Ala-scan (Figure 6 A) showed that α-syn binding is particularly sensitive to mutations in the C-terminal part from R11 onwards and to a lesser extent to mutations in the N-terminus. Truncation studies (Figure 6B) confirmed this, with removal of the first 4 residues from the C-terminus (*i.e.* residues 13-16) leading to particularly low binding and a smaller effect from the N-terminus. Interestingly, binding was temporarily restored to wt levels after the first (or last) 4 residues were removed but then binding generally declined. However, the general composition of the peptide was also favourable to α-syn binding, since 20 scrambled versions of penetratin gave rise to an average binding intensity of 30 ± 5% (data not shown).

**Figure 6.**
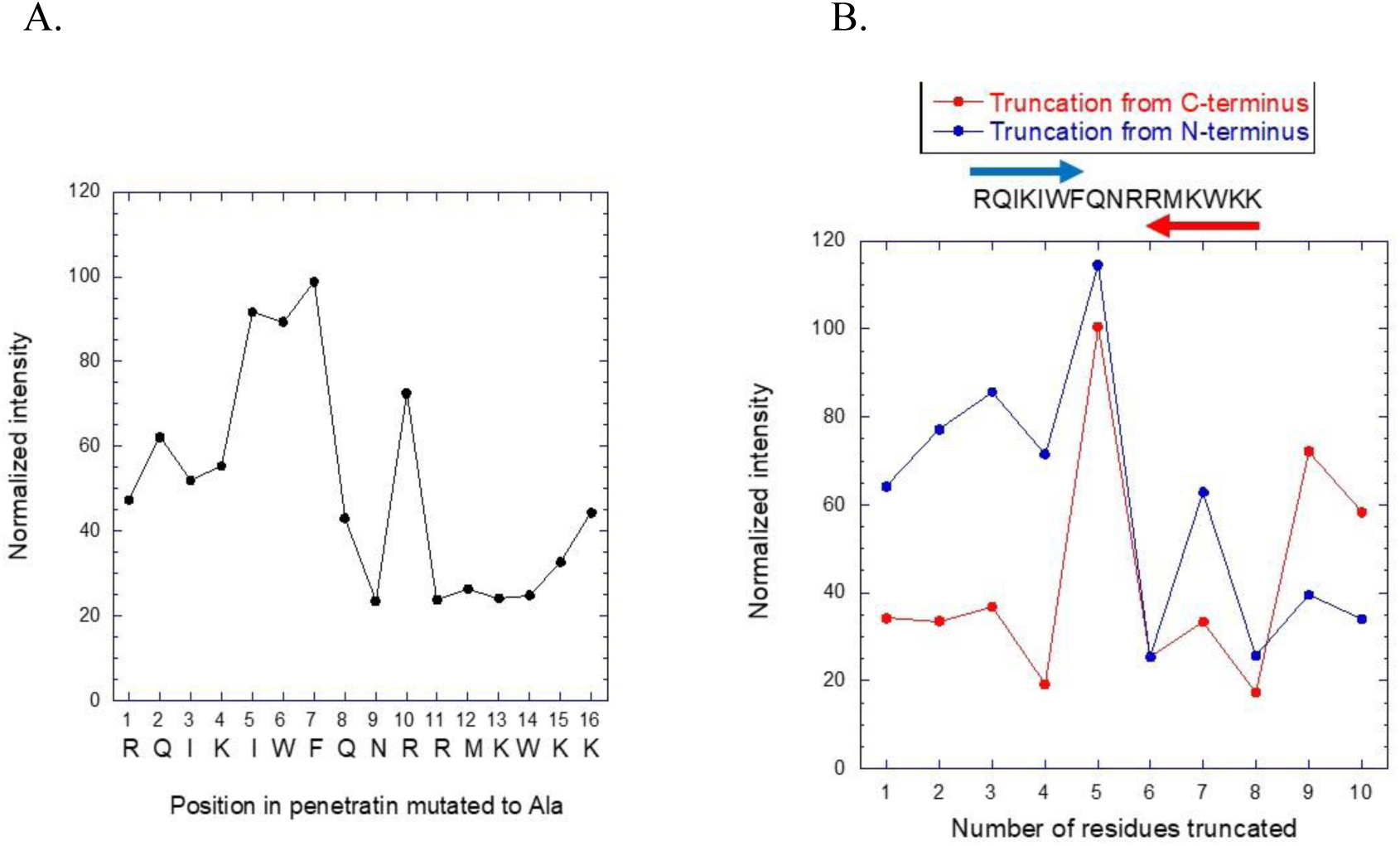
Binding of α-syn to penetratin measured by peptide arrays. Displayed peptides on the array were exposed to 0.05 mg/ml labelled α-syn and the binding was then quantified by densitometric scanning. Intensities are normalized to binding to wildtypepenetratin. (A) Ala-scanning of penetratin. (B) Terminal truncation of penetratin, leading to peptides down to 6 residues in length.

### Peptides efficiently decrease α-syn aggregation in *C.elegans* model of PD

The nematode worm *C. elegans* has been used extensively as a live animal model to gain understanding of PD. To examine the effect of the peptide on α-syn aggregation, we used a transgenic *C. elegans* strain (NL5901) overexpressing human α-syn-GFP in body wall muscles under control of unc-54 promoter. Worms were raised onto growth media containing *E. coli* OP50 supplemented with scrambled peptide, penetratin or cyclic-penetratin peptide. For fluorescence imaging studies, anterior portion of the worms were selected and kept constant across all groups, followed by quantification of GFP fluorescence which is represented in arbitrary units (Figure 7 A-E). A decrease from higher to lower fluorescence units reflects decrease in the GFP tagged α-syn aggregation. Compared to the scrambled peptide, there was a significant decrease in fluorescence levels upon worm treatment by both penetratin (45% decrease) and cyclic-penetratin (43% decrease) (Figure 7 E), suggesting that both peptides inhibited α-syn aggregation.

**Figure 7.**
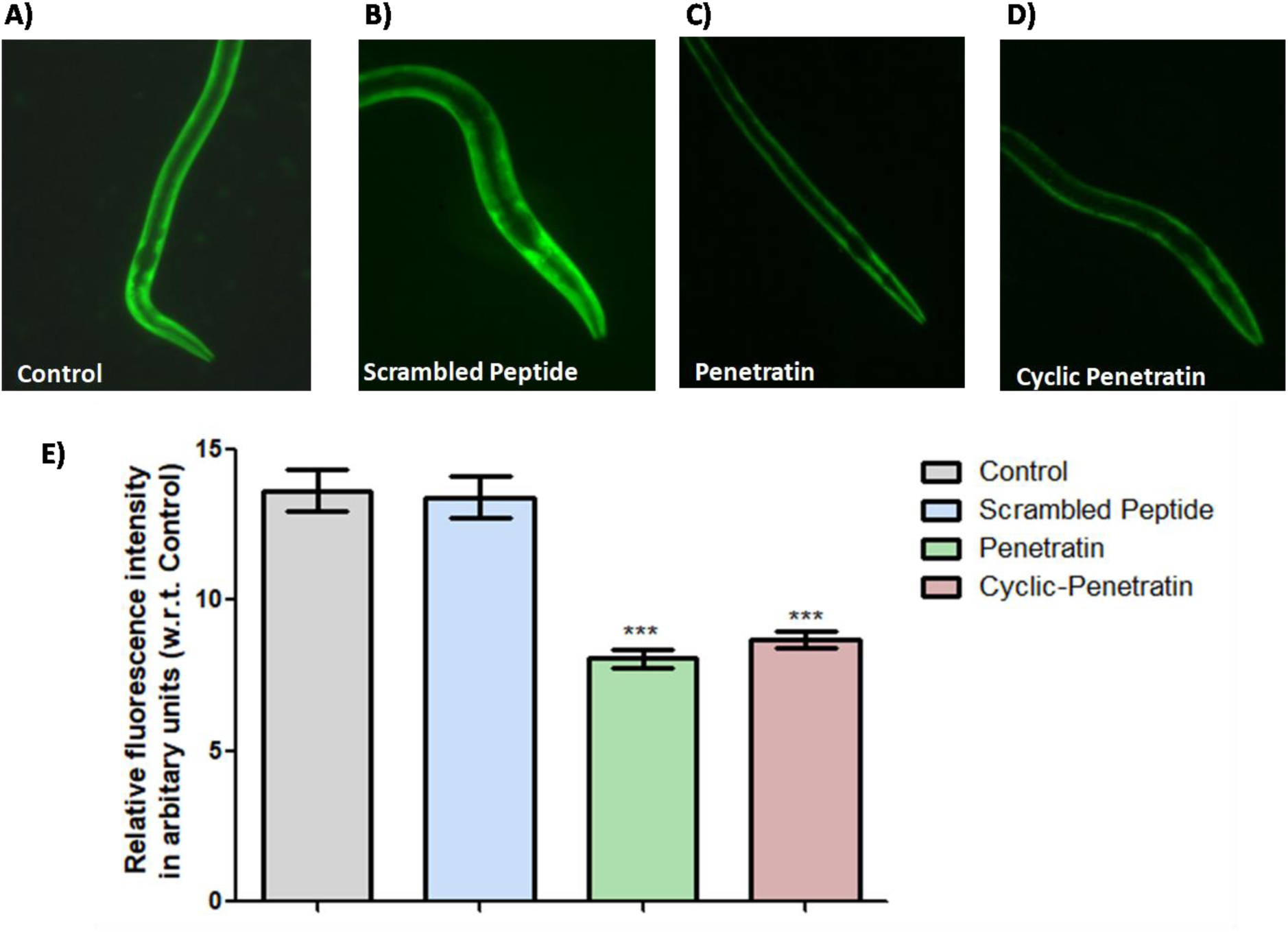
α-Synuclein aggregation in NL5901 strain of *C. elegans*. **(A-D)** Worms treated with OP50 and with, control, scrambled, penetratin or cyclic-penetratin. **(E)** The mean fluorescence intensity (indicative of α-syn aggregation) upon treatment of worms with indicated peptides. The fluorescence intensity was quantified using Image J software. As seen the penetratin and Cyclic Penetratin decrease α-Synuclein aggregation as compared to scrambled peptide. Statistical significance is as *p<0.05, **p<0.01, ***p<0.001. Scale bar, 50 µm.

### The peptides show potential improvement in overall dopamine health and rescue aggregation associated effects

Next, to visualize dopaminergic neurons, the *C. elegans* strain BY250 was examined using fluorescence microscopy. *C. elegans* has four bilaterally symmetric pairs of dopaminergic (DA) neurons that include two pairs of cephalic (CEP) neurons and one pair of anterior deirid (ADE) neurons (located within head region) and another pair is that of posterior deirid (PDE) neurons (located in a posterior lateral position).The strain BY250 expresses GFP under the influence of dopaminergic promoter (Dat-1), which is a transporter of dopamine. Relative fluorescence intensity of Dat-1 was measured in arbitrary units in the anterior region across various groups and analyzed in comparison to the control group (Figure 8A and 8B). Soma along with neurites in the head of penetratin-treated worms exhibited the highest fluorescence with an increase of 1.36 fold, followed by cyclic-penetratin with 1.18 fold increase (Figure 8B).

**Figure 8.**
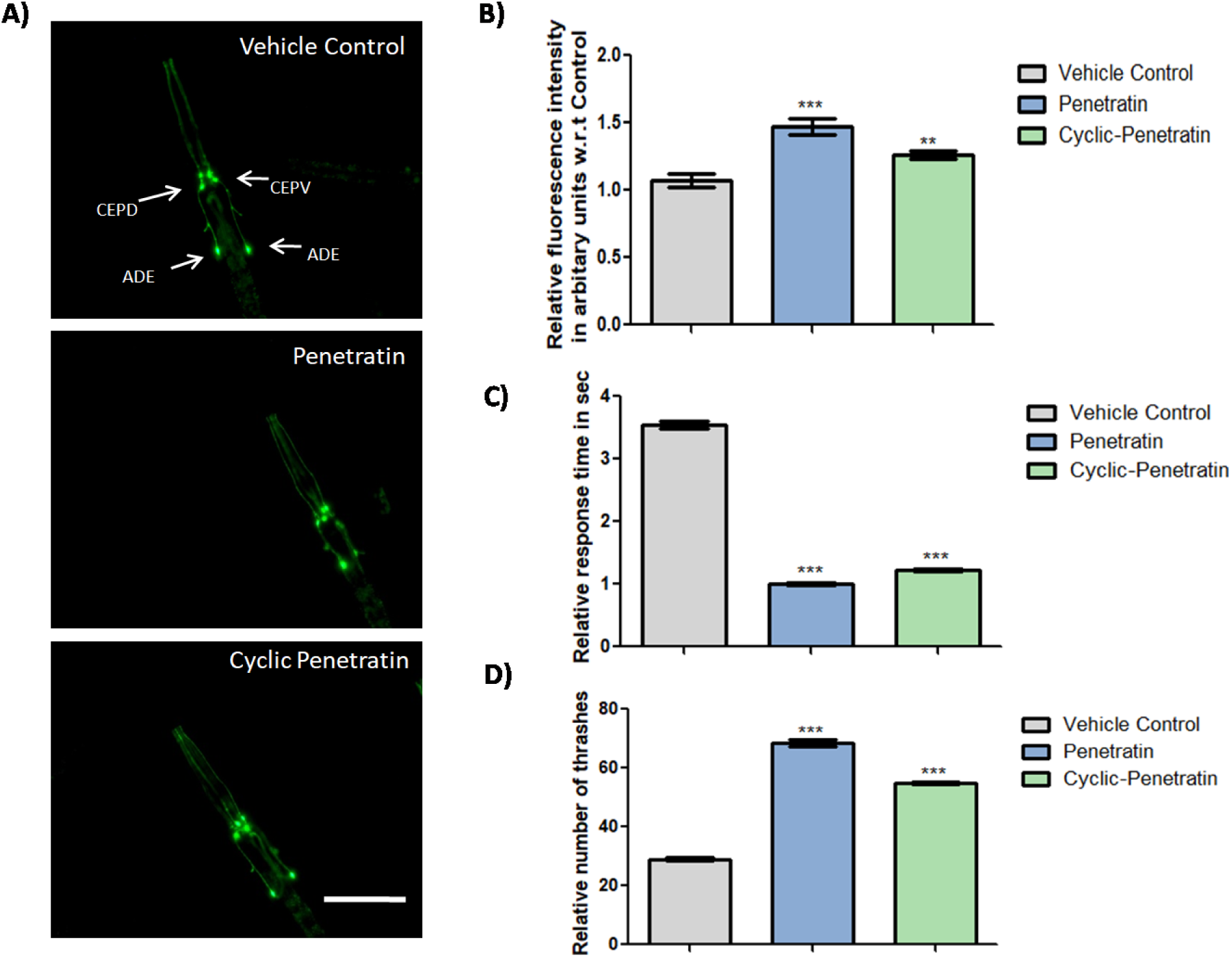
Effects of peptides on the end points associated with α-Synuclein aggregation and overall dopamine health. **(A)**Effect of peptides on expression of dopamine transporter dat-1 employing transgenic *C.elegans* strain BY250 (P*_dat-1_*::GFP) . Worms were raised on vehicle control(1x PBS+OP50 feed), 1% working concentration of penetratin and cyc- penetratin in vehicle 1X PBS. The images were quantified using ImageJ software. Scale bar, 50 µm.**(B)**Representative graph of the quantified GFP expression from A. As seen there is increased fluorescence in the soma and neurites of the DA neurons in the head region. As compared to the control,penetratin treated worms show highest fluorescence(per group n=10). **(C)**To monitor impact of peptides on worm’s dopamenergic health, nonanol repulsion assaywas carried out in transgenic strain of *C. elegans*NL5901 expressing α-syn. (Response time is inversely proportional to healthy function of dopaminergic neurons; lower the response time, healthier the dopaminergic function). (per group n=10).**(D)**Thrashing assay for motility reveals that peptide treatment led to increased number of thrashes per minute, which happens as a result of better motility (per group n=5). The graphical data was plotted using Non parametric student’s t-test using GraphPad software where *p <0.05,**p < 0.01,***p <0.001 significance respectively.

Further, to study the effect on dopamine health, we carried out the well-established 1-nonanol assay that estimates the changes in dopamine signalling in worms due to changes in dopamine level. Healthy worms with normal dopamine function will have an immediate repulsion response while worms with altered dopamine function would show a delayed repulsion response. The nonanol assay was carried out with NL5901 strain, in the presence or absence of peptides as described in materials and methods section. As seen in Figure 8C, the mean repulsion time for the α-syn transgenic worms (NL5901) was observed to be 3.5 sec. Interestingly, worms treated with penetratin or cyclic-penetratin showed significantly lower mean repulsion time of 1.0 sec (decrease of 0.71 fold) and 1.2 sec (decrease of 0.66 fold) respectively, suggesting that these peptides lead to restoration of dopamine signalling in worms.

Mobility and locomotion are very finely regulated functions in *C. elegans,* as worms exhibit a typical sinusoidal wave pattern while moving. This requires a fine balance of excitatory and inhibitory neurotransmission, which if altered by any chemical, genetic or physiological event, will lead to its disruption. This is reflected in a very well established “thrashing assay”, wherein worms are put in a drop of buffer followed by observation of their natural thrashing response and any disruption thereof. We hence subjected the peptide treated NL5901 worms to thrashing assay; the numbers of thrashes per 30 seconds were counted and the recorded number of thrashes were plotted. We observed that, in the untreated group the mean number of thrashes observed were 35 per 30s, while worms treated with penetratin and cyclic- penetratin exhibited a thrashing count of 68 and 55 per 30s, respectively. Hence, in the performed experiment it was clear that locomotion is significantly increased after treatment with the peptides in the NL5901 strain that overexpress α-syn (Figure 8D).

### Penetratin reduces the decline of locomotor coordination in PD mice

The increased death of dopaminergic cells in PD affects motor coordination ability. We thus examined whether penetratin and cyclic-penetratin that reduced α- syn aggregation, and improved dopaminergic cellular health in *C. elegans* could also improve locomotor coordination in aging PD mice. The transgenic mice expressing A53T mutant of human α-syn under Thy1 promoter were used as PD model. The mouse model expresses α-syn-A53T in the brain regions and show typical motor symptoms as observed in clinical PD. The transgenic PD mice were subcutaneously treated with saline (placebo control, PC) or 30/60mg/kg body weight of penetratin, every 3^rd^ day, starting from 12 weeks of age and continued until 22 weeks. The non-transgenic mice (WT) were used as controls with respect to the PC group for comparison. Penetratin was administered subcutaneously to PD animals and motor coordination activity was monitored using rotarod and wire hanging tests. The muscular strength and coordination of all the animals in four groups were evaluated in terms of fall off time, acceleration and distance travelled by the animals on a rotating rod. As seen in Figure 9A, there was no significant change in the fall-off time in WT animals from 12-22 weeks of age. The mean fall-off time from the accelerating rotating rod decreased significantly for transgenic A53T mice (PC) with age (Week 12:Week-18:Week-24::25sec:11sec:7sec respectively). Interestingly, as compared to mice in PC group, relatively less decrease in falling-off time from the rotating rod was observed in A53T mice administered with lower dose (30mg/kg) of penetratin. The animals treated with 60mg/kg of penetratin also showed significantly improved latency in falling off the rotating rod (increased by ∼47%) on the 22^nd^ week, as compared to untreated A53T mice (PC).

**Figure 9:**
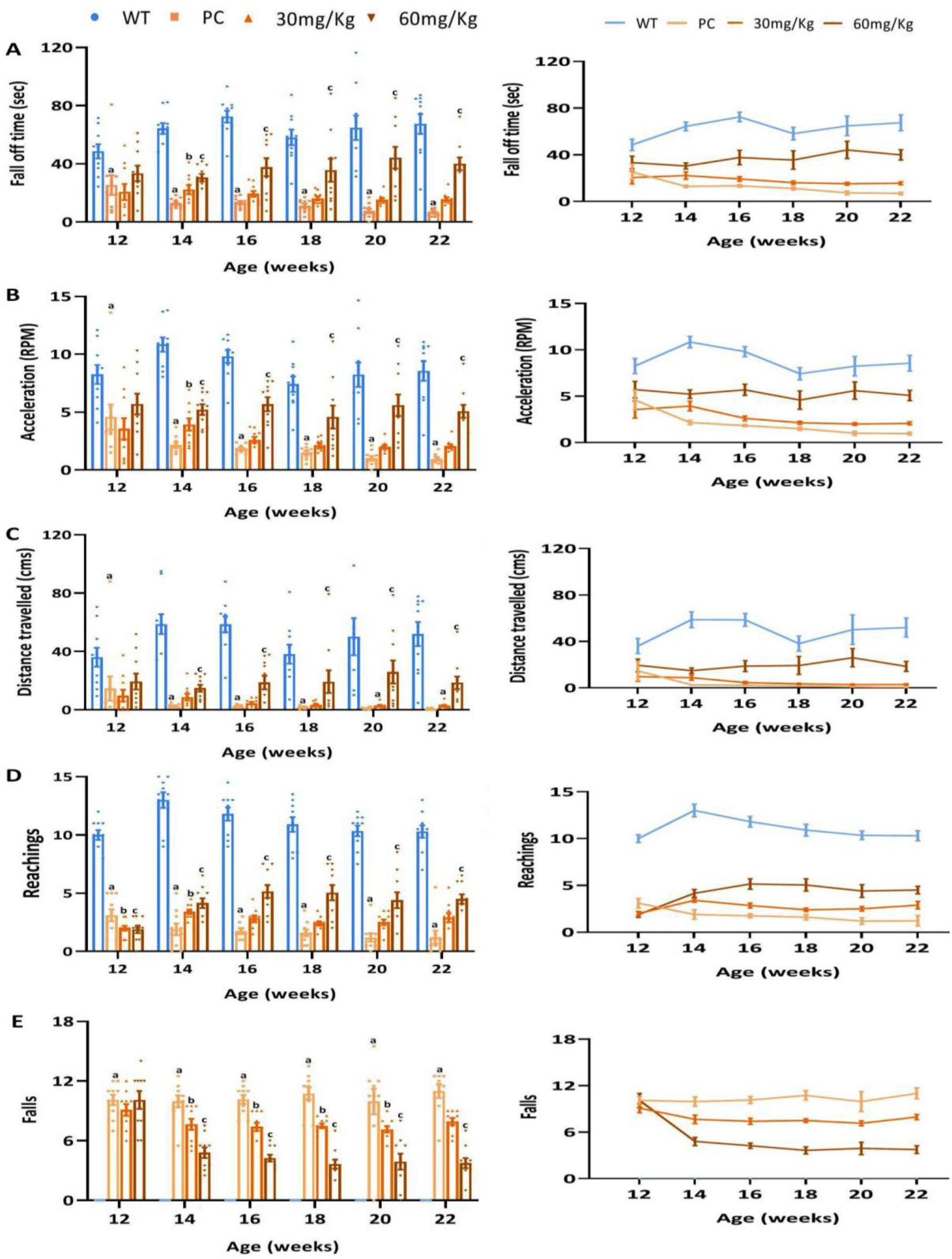
Effect of Penetratin treatment on locomotor coordination activity assessed using rotarod and wire hanging test in transgenic PD mice. The PD mice were treated with indicated amount of penetratin, and rotarod (A-C) orwirehanging (D-E) test were carried out to assess neurobehavioral functions. (A-C) The WT or PD mice (with or without (placebo control) penetratin treatment) were placed on rotating rod, and their latency to fall (A), ability to withstand acceleration (B) and distance travelled (C) were recorded. (D-E) The animals were placed onto a hanging wire and their ability to reach opposite end (D) or fall from wire were recorded. PC shows the placebo group. Values are expressed as mean±SEM, n=10. P˂0.05, **^a^**significantly different from WT group, **^b, c^**significantly different from PC group. The mean value of the data obtained is represented as bar graph (left panel) and also as line curve (right panel) for more clarity.

We further examined motor impairments in penetratin treated and untreated mice with respect to their ability to withstand acceleration and distance travelled on the rotating rod. The mice treated with 30mg/kg (peptide/mice weight) of penetratin showed significant improvement in their ability to withstand acceleration (Table S2). The A53T mice treated with higher dose (60mg/kg) of peptide could significantly withstand higher acceleration of the rotating rod as compared to PC or 30mg/kg peptide treated animals (Figure 9B). In addition, there was a significant reduction in the distance travelled by PC mice with age; however the distance travelled by the penetratin treated mice (in 60mg/kg treated group) on the 22^nd^ week remained near to that observed on the 12^th^ week (Figure 9C), suggesting a beneficial therapeutic efficacy of peptide in managing locomotor coordination in A53T mice.

Similar to as observed in rotarod test, we observed significant improvement in penetratin treated versus PC mice on wire-hang test (Table S3). For PC mice, number of reachings decreased from ∼3 (12^th^ week) to ∼1 (22^nd^ week) (Figure 9D). For A53T mice administered with penetratin at 60mg/kg, the number of reachings increased from ∼2 (12^th^ week) to ∼4-5 (22^nd^ week). Similarly, the number of falls in 60mg/kg penetratin treated mice decreased from ∼10 (12^th^ week) to ∼4 (22^nd^ week) (Figure 9E).

### Penetratin restores histological imperfections in PD mice

The mice in all the groups (with and without penetratin treatment) were sacrificed for assessing α-syn mediated histopathological changes in cortex of mice by Haematoxylin and Eosin (H&E) and Congo red staining. Histological evaluation through H&E staining showed that wt mice (control group) possess normal tissue morphology (Fig. 10 A-D). Whereas, brain sections from PD mice without treatment (PC) showed noticeable degeneration in neurons, wherein cells appeared to be pyknotic and damaged, accompanied by condensed and increased void space around the neurons, thus validating degeneration in the affected tissue region. However, sections from mice treated with 60 mg/Kg of penetratin for 22 weeks showed more number of neuronal population accompanied by improved morphology with respect to those treated with 30 mg/Kg of the peptide and PC mice.

**Figure 10.**
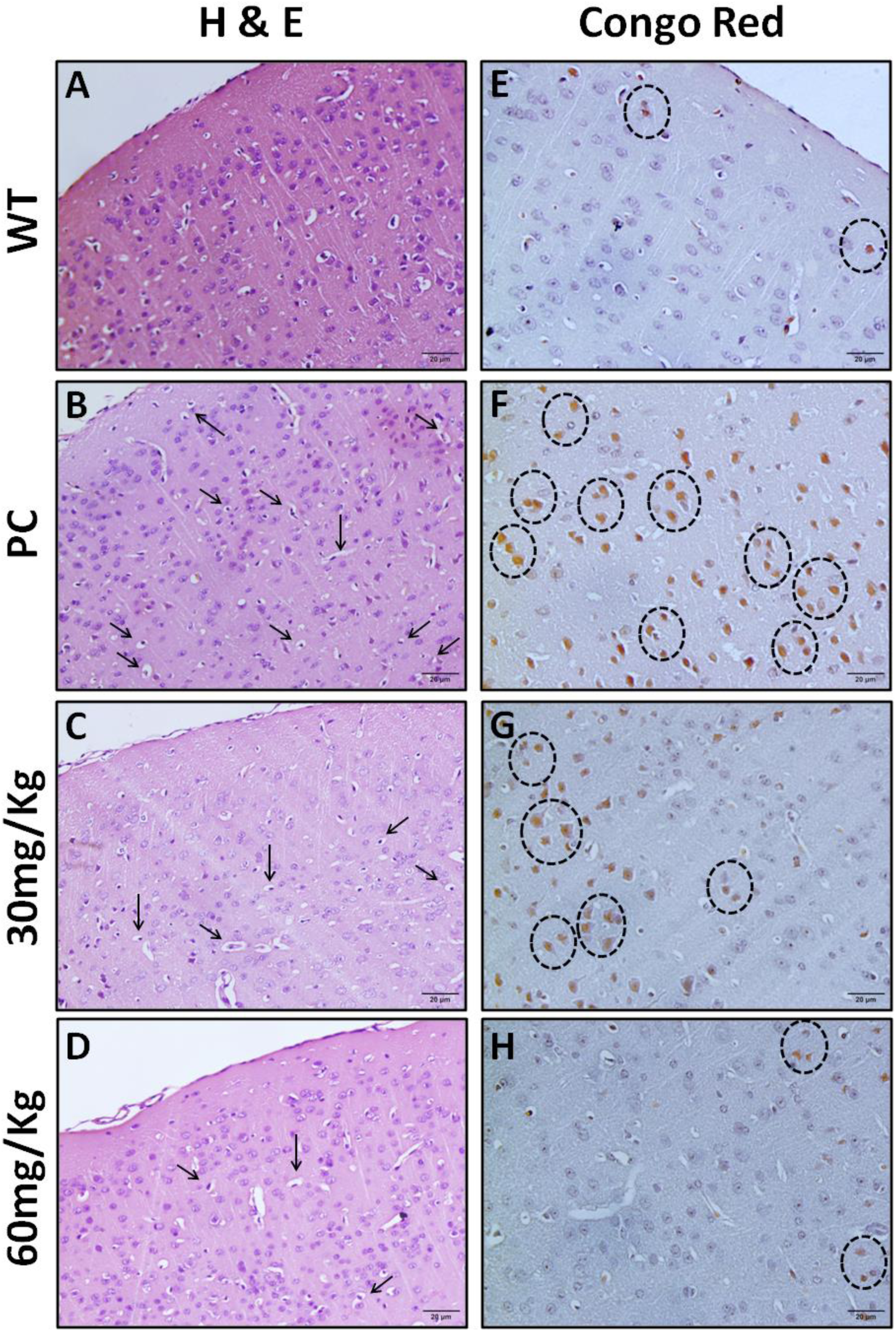
Effect of penetratin treatment on histological changes observed using H&E staining (A-D) and Congo red staining (E-H) in cortex tissue sections from mice brain. Magnification 40× (scale bar-20µm).

In addition, Congo red staining showed analogous morphological features as observed in H&E staining (Fig. 10 E-H). Examination of Congo red stained sections in PC group showed higher number of orange colored amyloid deposits within the cortical layer, along with appearance of large vacuolated and pyknotic nuclei in neuronal cells. Whereas, cortical section from Wt controls and PD mice treated with 60 mg/Kg of peptide showed significantly reduced number of amyloid deposits and less pyknotic nuclei with respect to those treated with 30mg/Kg of penetratin or with PC alone. Thus, histological examination confirmed that treatment with 60mg/Kg of penetratin for 22 weeks is effective in maintaining neuronal morphology in PD mice.

## Discussion

Despite extensive efforts, there is currently no disease-modifying therapy for managing PD. The current therapies are aimed at restoring dopamine levels which improves motor functions, however they have limited or no beneficial effect on non-motor symptoms and also cause undesired side effects. Though various genetic risk factors such as α-syn, PINK1, LRRK2, PARK1 and GBA1 are associated with PD, neuronal accumulation of α-syn aggregates is the most commonly observed pathological feature observed in PD (Franco et al., 2021). Thus, among various approaches, those that target α-syn aggregation continue to be an extensively explored therapeutic intervention. In the present study, we demonstrate that penetratin is a potent inhibitor of α-syn fibrillation, and proves its therapeutic efficacy in *C.elegans* and mice model of PD.

As α-syn aggregates primarily accumulate in the cytoplasm, the ability of a potent inhibitor/drug to traverse through cellular membranes and gain access to α-syn and its aggregates is crucial for determining its efficacy. Thus, we screened various CPPs that are well known to permeabilize through cellular membranes. Such peptides have long been used to deliver cargos into the cell cytoplasm. Among the cationic and amphipathic classes of examined peptides, we did not observe any correlation between their physicochemical properties and ability to inhibit α-syn fibrillation. Though IMT-P8, MAP and R8 promoted fibrillation, TP-10, SynB3 and penetratin inhibited it. Among all peptides that were screened, penetratin was found to be the most potent inhibitor of α-syn aggregation.

The α-synis knownto undergo conformational transition from its disordered state to β-sheet rich structure formation. Such conformational transition is known to be a crucial step in the conversion of disordered proteins into amyloid fibrils.In molecular docking simulations study, we observed temperature-induced transitions of intrinsically disordered α-syn to a β-rich metastable state. Similar high β-content metastable states that can propagate to fibrillary states, have previously been identified upon temperature induced misfolding of prion protein(Chamachi and Chakrabarty, 2017). Interestingly, pre-incubation with penetratin inhibited these conformational changes into a β-sheet rich state. The inhibition of these transitions by peptides could be attributed to H-bond and salt bridge interactions with non-amyloid-β-component (NAC) regions of α-syn (Table S4 and S5). These peptide-protein interactions can interfere with forces driving transitions of intrinsically disordered α-syn to a β-sheet rich metastable state.

As compared to their linear counterparts, cyclic derivatives are relatively more stable, less sensitive to both exopeptidases (due to cyclization of termini) and endopeptidases (due to increased conformational rigidity). The penetratin cyclized through N- and C-termini also inhibited α-syn fibrillation, suggesting that structural constraints introduced through cyclization do not have an adverse effect on peptide activity. The binding study showed that the cyclic derivative binds with a relatively lower affinity to α-syn presumably due to increased rigidity of the cyclic-penetratin.

The region 61-95, known as NAC, forms the amyloid core upon α-syn aggregation (Li et al., 2018; Tuttle et al., 2016a). Intriguingly, our MD simulation studies demonstrate that penetratin primarily interacts with the C-terminus of α-syn due to electrostatic interactions among the negatively charged C-terminus and positively charged residues present in the peptide. NMR studies further confirmed these interactions of linear penetratin with C-terminus of α-syn. Similar to penetratin, the cyclic-penetratin peptide was also found to interact with the C-terminus of the protein suggesting that cyclization has no effect on the nature of interaction between the peptide and α-syn. Previous studies show that the C-terminus plays an important role in formation of α-syn fibrils by interacting with the NAC region of the protein. It is possible that the interaction of the peptide with α-syn inhibits the C-terminus from interacting with the NAC region, thereby preventing the protein from undergoing β- sheet rich state transition.

The decrease in the α-syn aggregation in *C.elegans* upon peptide treatment (penetratin and cyclic-penetratin) further corroborates the *in-vitro* findings that the peptides are potent inhibitors of α-syn fibrillation *in-vivo*. Further, increase in dopamine health measured using nonanol and thrashing assays shows that peptides inhibit α-syn aggregation and its associated toxicity. This is also in agreement with our finding observed with SH-SY5Y cells, wherein fibrillation reactions containing α- syn and either peptide were found to be less toxic as compared to α-syn fibrils alone. PD is an age-related disorder in which the affected regions of brain degenerate with age resulting in progressive development of the disease symptoms such as motor incoordination. The increase in fall-off time, decreased travel distance, with concomitant increased number of falls in PC mice during various motor coordination tests are consistent with neurodegenerative effects as observed in PD. Interestingly, treatment of penetratin prevented the further deterioration in motor coordination skills of PD mice suggesting *in-vivo* efficacy of penetratin. Moreover, histological imperfections in PC mice were found to be reversed upon treatment with 60mg/Kg of penetratin for 22 weeks. Collectively, our study demonstrates the efficacy of penetratin against α-syn associated toxicity in multiple PD models, however additional clinical investigations are warranted to further validate the use of penetratin in treating PD.

With a change in lifestyle and an increase in life expectancy, the neurodegenerative diseases are becoming more prevalent. The disease not only affects the patient but also the caregivers as the patient becomes extremely frail and requires constant care. Interestingly, the underlying cause of many such diseases is the neuronal accumulation of morphologically similar aggregates which results in neuronal toxicity. The similarity of the underlying basis suggests the possibility of common therapeutic agents against many of these diseases. The present finding, that penetratin inhibits α-syn fibrillation and imparts therapeutic effects in pre-clinical models of PD, opens avenues for the development of similar lead molecules against a number of neurodegenerative diseases.

## Acknowledgements

We acknowledge the support of Mr. Ashish Dubey, Dr. Sachin Raut, Mr. Bhupinder Singh, and Ms. Madhumita Dey for initial mice experiments. We also acknowledge support from Rashami Rajwansh for initial *C. elegans* study.

## Materials and Methods

### α-syn purification

For α-Syn protein expression, pT7-7 based expression plasmid was transformed into *E. coli* Rosetta (DE3). Expression of α-Syn protein was induced using 0.5mM IPTG at 30°C. After 8 hrs, cells were harvested and pellet was resuspended in10 mMTris-HCl, pH 8.0, containing1 mM EDTA and 1 mM protease inhibitor cocktail (Thermo Pierce). Further, Protein purification was performed using method described before with slight modifications (van Raaij et al., 2006). Briefly, cells were lysed, and cellular lysate was boiled at 95°C for 30 minutes. The α-syn was precipitated by adding 50 percent ammonium sulphate, then incubating for 1 hour on ice with regular shaking every 10 minutes. The pellet was separated and washed with equal amounts of 100 mM ammonium acetate and 100% ethanol. The pellet was then dissolved in 10 mM HEPES, pH 7.4 containing 50 mM NaCl, and dialyzed extensively to remove excess ammonium sulphate. Protein purity was confirmed on 15% SDS-PAGE.

For NMR experiments, ^15^N-labelled α-Syn protein was expressed in *E. coli* BL21 (DE3) and purified with slight modification (Marley et al., 2001). Briefly,BL21 (DE3) cells transformed with plasmid encoding α-Syn were grown in M9 minimal media supplemented with ^15^NH_4_Cl as the sole nitrogen source and induced for four hours by IPTG at 0.8 O.D.600nm. The bacterial cells were harvested and ^15^N- labelled α-Syn protein was purified as mentioned above.

### Fibrillation Kinetics

400 μM of purified α-syn containing 4 mM ThioflavinT (ThT) with and without peptides at equimolar concentrations was added in a 96 microwell flat bottom black plate to a final volume of 100μl. The fibrillation reaction was then initiated at 37°C in a fluorescence plate reader with a800 rpm shaking speed. The ThT fluorescence intensity was monitored at regular interval of time upon excitation and emission at 442 nm and 482 nm respectively.

### Transmission Electron Microscopy

The morphology of the oligomeric species was assessed using transmission electron microscope (TEM) JEM-2100 (JEOL, Japan). For TEM imaging, samples were adsorbed onto 300 mesh carbon-coated grids following by staining with 1% Phosphotungstate for 30 seconds, followed by visualization under the electron microscope Images were captured using JOEL software and analyzed.

### Confocal microscopy

For examining α-syn-GFP punctae in SY246 yeast strain, cells were induced with galactose for 12h and further incubated with and without penetratin or cyc- penetratin (200µM each) for 60 minutes. Later, cells were mounted on 1% agarose blocks for imaging. The α-syn-GFP was excited at 488nm using argon laser in confocal microscope (A1 plus Ti ,Nikon, Japan) and images were recorded using NIS elements software. Approximately, 200 yeast cells showing α-syn-GFP expression were analyzed.

### SDS-PAGE analysis

For SDS-PAGE analysis, samples were taken before (0h) and after fibrillation (7h), and fractionated by centrifugation at 4000 rpm for 1 minute into supernatant and pellet. Each fraction was re-suspended into SDS loading dye and further separated on 15% SDS-PAGE. The fractionated proteins were seen on SDS-PAGE using Coomassie Brilliant Blue G-250 staining.

### CD spectroscopy

Samples were collected from the saturation phase of fibrillation reactions. The reactions were diluted to 5 µM (α-syn). The CD spectra were recorded for the reaction mixture and also of the supernatant obtained after centrifugation at 4000 rpm for 1 minute. Far-UV CD spectra were monitored at a scan rate of 10nm/min over a range of 250-195nm in JASCO-J-815 spectropolarimeter (JASCO, MD, USA)in a cuvette of 1-mm path length. The Mean Residue ellipticity (Ɵ_MRE_) was calculated from raw data as follows:

Ɵ_MRE_= (100*Ɵ_obs_)/[d*C*(n-1)]

Where Ɵ denotes the observed ellipticity (in degrees), d the path length (in centimeters), C as protein concentration (molar), and n, the number of amino acids in the protein. Here, Ɵ_MRE_ of supernatant was subtracted from Ɵ_MRE_ from total reaction mixture to get the CD spectra for fibrils.

### Peptide synthesis

Peptides were synthesized at a peptide synthesizing facility at CSIR-Institute of Microbial Technology (CSIR-IMTECH), Chandigarh, India and/or purchased from Genscript Biotech Corporation (New Jersey, USA). Briefly, these peptides were synthesized by solid phase peptide synthesis using Fmoc (N-(9-fluronyl)- methoxycarbonyl) chemistry (Gautam et al., 2015a).

### Microscale thermophoresis studies

MST experiments were performed with labelled α-syn using Monolith NT™ Protein Labelling Kit RED-NHS (NanoTemper Technologies). To perform thermophoresis, penetratin and cyclic-penetratin peptides were serially diluted from 100 µM–3.05 nM keeping α-syn concentration constant at 40nM in HEPES buffer (pH 7.4). After equilibration, the reactions were loaded into Monolith™ NT.115 standard capillaries for thermophoresis. Data was analyzed using MO affinity analysis software. The normalized values of dose response (ΔF_norm)_ obtained for each concentration of peptides (penetratin or cyclic-penetratin) was plotted against the ligand concentration. Data was fitted to obtain the dissociation constant. The results are average of duplicate values with standard deviation (S.D.) from two different experiments.

### MTT Assay

MTT experiments were performed in human neuroblastoma SH-SY5Y cells to assess α-syn mediated cellular toxicity. SHSY5Y cells were maintained in DMEM/F12 media (Gibco) with 10% heat-inactivated fetal bovine serum (Gibco) with a mixture of 0.1% penicillin+streptomycin (Gibco) antibiotics. About 10,000 cells were seeded in triplicates in a sterile 96-well plate. A total of 100 μL of growth media mixed with sonicated fibrillation reaction mixture of 10μM α-syn alone or with peptides was added into the cells. After 24 h of treatment at 37°C in the presence of 5% CO_2_, MTT (10µM, 5 mg/ml) was added and again incubated for 4h at 37°C. Further, 100 μL of DMSO was added and the formed formazan crystals were dissolved by gentle pipetting, followed by 4 h of incubation at 37°C in CO_2_ incubator. The absorbance was monitored at 570 nm in a multimode plate reader.

### Molecular dynamics system setup and simulations

*Simulation setup:* All simulation experiments were conducted using the GROMACS 5.1 software suite with the gromos 54A7 force field (Huang et al., 2011; Lin and van Gunsteren, 2013; Schmidt et al., 2011). Prior to testing the impacts of various biomolecules, the human micelle-bound α-syn (1xq8) structure was pre-simulated at 473 K to obtain random protein conformation (as found during CD tests). Two different types of systems were set up to explore the interference of intercalators with amyloidogenic pathways: simulations of α-syn in the absence and presence of penetratin. In the first system, pre-generated random conformation penetratin monomers and protein were simulated for 100 nanoseconds, with penetratin monomers put in random orientation in individual cubical boxes and protein in the centre, according to their experimental aggregation conditions.

For protofibril simulations, only α-syn fibrils (PDB id: 2N0A) were employed, with the fibril assembly alone placed in the centre of a box with the same dimensions as before (Tuttle et al., 2016b). Following that, penetratin was placed in the box at random in each arrangement. Each system was specifically solvated with a sufficient amount of SPC water molecules and ions for electroneutrality at a concentration of 0.15 M NaCl, which corresponded to previous experimental settings. After that, equilibration was done using an isothermal-isobaric and isochoric-isothermal ensemble for 1 ns each. Long-range electrostatics with short-range interactions were treated using PME at a cutoff radius of 10Å for both coulomb and van der Waal potentials. Using a velocity rescale and a Berendsen barostat, the system temperature and pressure were kept constant at 310K and 1 bar respectively. Finally, the velocities were calculated using a Leapfrog integrator to solve Newton’s equations of motion with a 2 fs time step. At each 5 ps interval, the coordinates were recorded. The images were produced using the PyMol package, and the resulting trajectories were analyzed using the inbuilt gromacs capabilities.

### Preparation of labelled low molecular weight (LMW) α-Syn for NMR experiment

The buffer was exchanged with phosphate buffer (PB) (20mM), 50 mMNaCl (pH 6.6) by passing through a 3 kDa cut-off filter (MWCO, Millipore). Further, the protein solution was passed through pre-washed centricon YM-100 filter (100 kDa MWCO, Millipore) and centrifuged at 10,000 g for 20 min at 4°C and the flow through containing α-Syn<100kDa (LMW α-Syn) was used for the NMR experiments. 5 mM stock of linear and cyclic-penetratin was prepared by dissolving the peptides in the 20 mM PB with 50 mM NaCl (pH 6.6) for the NMR experiments.

### Nuclear Magnetic Resonance

Two-dimensional ^1^H-^15^N heteronuclear single quantum coherence spectroscopy (HSQC) was captured for 150 µM ^15^N labeled α-Syn in 20 mM PB containing 50 mMNaCl (pH 6.6) at 288 K on Bruker Ascend 750 MHz spectrometer equipped with 5 mm triple resonance TXI probe. Series of ^1^H-^15^N HSQC was acquired at various increasing molar equivalents (0.2, 0.4, 0.6, 0.8, 1.0, 2.0, 5.0, and 8.0) of penetratin and cyclic-penetratin. The spectra were acquired for 8 scans with 256 and 2046 data points in the direct and indirect dimensions. All the spectra were recorded on Bruker Topspin 3.5pl6 software and analyzed by CcpNMR (Collaborative Computational Project for NMR) software. The α-Syn amide cross-peaks were assigned by the ^1^H-^15^N chemical shift values from BMRB and reduced dimensionality experiments.(Ref) The intensities of each amino acid amide cross-peaks were derived from CcpNMR software. The intensities of amide cross-peaks of α-Syn in presence of different molar equivalents of peptides were normalized with that of in absence of peptides. Chemical shift perturbation (CSP) was calculated by using:

CSP=√(δH^2^+(0.1δN)^2^)

where, δH and δN represent the difference in the chemical shifts of proton and nitrogen of amide cross-peaks of α-Syn in absence and presence of different molar equivalents of peptides respectively. 3KD was derived from the NMR titration by the following equation:

Δδ_obs_ = Δδ_max_ {([P]_t_ + [L]_t_ + K_D_) – [ ([P]_t_ + [L]_t_ + KD)_2_ - 4[P]_t_ [L]_t_ ]^1/2^} / 2[P]_t_

where, Δδ_obs_ is observed CSP at a particular concentration of ligand, Δδ_max_ is maximum shift change upon saturation, [P]_t_ and [L]_t_ is total protein and ligand concentration respectively, and KD is dissociation constant.

### *C. elegans* culture and its maintenance

Experiments to study the effect of peptides in *C. elegans*, were carried out following the standard liquid culture protocol. Briefly, *C. elegans* nematodes were grown on S Medium added with *E. coli* OP50 as a food source. Further, age synchronized embryos after axenization were transferred to screening flasks. In this study, transgenic worms NL5901 {unc-54:: α-synuclein::YFP+unc-119} BY250 (Pdat- 1::GFP) and wild type Bristol N2 were used . Strain NL5901 and N2 were purchased from the Caenorhabditis Genetics Centre (University of Minnesota, USA) and strain BY250 was a kind gift from Prof. Randy Blakely (Florida Atlantic University Brain Institute, USA). All strains were maintained at 22°C and optimum conditions were followed for their propagation in liquid culture (Stiernagle, 1999).

### Primary screening of the test peptides for the effects on α-Syn aggregation in C. elegans

α-Syn aggregation being the hallmark of PD pathology, the NL5901 strain of C. elegans expressing human α-syn were employed to screen the test peptides towards reducing the α-Syn expression. Following treatment of worms with 1% test peptides as part of the liquid culture medium for 48 h, L4 stage synchronously staged worms were harvested, washed and subjected to fluorescence microscopy using Cral Zeiss AxioImager Z4 microscope equipped with GFP filter (at excitation wavelength of 488nm). Worms were given multiple washes with M9 buffer to clean any debris, and then immobilized with sodium azide before staging the capture. Worms fed with OP50 served as controls whereas worms fed with scrambled peptide were tested alongside test peptides, so as to have an internal technical control for the presence of similar amino acids within the medium, but with scrambled sequence. Images were quantified using image J software (NIH, Bethesda, USA), where the fluorescence of the control worms and peptide treated worms was measured from the selected pharynx region uniform across the groups (Kumar et al., 2018). The fluorescent intensity for each group was then graphically illustrated using GraphPad 8.0 Prism statistical software (San Diego, California, USA).

### Effect of peptides treatment on locomotion in *C. elegans*

To investigate the effects of Penetratin and cyclic-penetratin on motor movements of NL5901 worms, thrashing rates were measured. In this experiment, worms were cultured in presence of control and test peptides for 48 h followed by washing with M9 buffer. Further, a drop of M9 buffer with worms was placed on a glass slide and movement of the worms was recorded for 30s using stereozoom microscope (Nikon, Japan). Frequency of sigmoidal body bends were calculated where the worm that oscillates its head and/or tail to the one side was counted as one thrash (Haque et al., 2020; Kumar et al., 2018). The counted thrashes for n=5 worms per group was analyzed by student’s t-test and graphically represented using GraphPad Prism 8.0 statistical software (San Diego, California, USA).

### Screening of the test peptides for the effects on dopamine transporter (DAT-1) in *C. elegans*

Dopamine transporter (DAT-1) is of importance in studying PD hence transgenic strain BY250 was employed to study the effect of penetratin and cyclic- penetratin on DAT-1 transporter. The strain BY250 expresses GFP specifically in its 8 dopaminergic neurons. Following the treatment of worms with 1% of peptides for 48h, the age synchronized L4 staged worms were subjected to fluorescence imaging. These worms were washed with M9 buffer and immobilized with sodium azide before imaging(Jadiya et al., 2011). Images were quantified using image J software (NIH, MD, USA), where the fluorescence from dopaminergic neurons and neuronal portion of head region of the control worms and peptide treated worms was measured within selected area keeping uniform across all the groups. The fluorescent intensity was represented using GraphPad Prism 8.0 statistical software (San Diego, California, USA).

### Effect of peptides treatment on dopamine signaling in *C. elegans*

Nonanol repulsion assay was performed to evaluate the effect of test peptides on DA-related functions. In *C. elegans*, olfaction is governed by dopaminergic neurons and worms with reduced dopamine levels exhibit a delayed response to 1- nonanol. In this assay, treated worms were washed with M9 buffer and transferred to a sterile NGM agar plate without food. Further, a drop of the odor-based repellent 1- nonanol was placed near worm head, and the worms’ response time to the repellent was measured (Haque et al., 2020; Kumar et al., 2018). The response time for n=10 per group was analyzed by Student’s t-test and graphically illustrated using GraphPad Prism 8.0 statistical software (San Diego, California, USA).

### Statistical analysis

Values represented for *C.elegans* experiments were expressed as mean±SEM. Statistical analysis was carried out using Graph Pad prism 5 software. Statistical significance between various groups was calculated using Student’s t test.

### Animals and treatment schedule

Transgenic mice [B6.Cg-Tg(THY1-SNCA*A53T)F53Sud/J] were obtained in breeding pairs from Jackson Laboratories (Bar Harbor, ME, USA) and housed at the animal care facility of the institute (iCARE at CSIR-IMTECH, Chandigarh, India). The pairs were used to generate a stable breeding colony of transgenic mice with mutation (A53T) in human SNCA gene, expressed through human THY1 (thymus cell antigen-1, theta) promoter. The identified mice hemizygous for the mutation were bred on a mixed background to produce transgenic and non-transgenic mice. To identify transgenic mice, tail samples from all the pups after weaning were collected and processed for DNA isolation using DNA isolation kit as per the manufacturer instructions (Qiagen). PCR amplifications were performed on isolated DNA samples according to the genotyping protocol provided by Jackson Laboratories (Bar Harbor, ME, USA).

Later, transgenic and non-transgenic male mice were identified (12 weeks of age, weighing ∼18-20g) and obtained from the animal facility for initially habituating them to the home environment for 4-7 days. Experimental procedures in mice were carried out between 09:00 and 15:00 h in accordance with the guidelines for ethical use and care of laboratory animals, following approval by the Institutional Animal Ethics Committee (IAEC/2020/2019). Non-transgenic mice were housed separately and served as controls (WT, 10 mice), whereas transgenic mice were divided into three groups (with 10 mice each) as follows:

WT (Control): Animals were non-transgenic and given vehicle alone.

Placebo control (PC): Animals were transgenic (THY1-SNCA*A53T) and given vehicle alone.

.PC + 30 mg/Kg treated:PC mice were treated with linear peptide at a dosage of 30 mg/kg body weight, every 3^rd^ day (subcutaneously) for 22 weeks.

PC + 60 mg/Kg treated:PC animals were treated with linear peptide at a dosage of 60 mg/kg body weight, every 3^rd^ day (subcutaneously) for 22 weeks.

### Analysis of behavioral functions

Behavioral functions in mice were assessed for evaluating locomotor coordination and muscular strength in following neurobehavioral tasks:

### Rotarod test

Mice were placed on a revolving rod that moves at a steady or accelerated pace to assess locomotor coordination in animals. The automated equipment keeps track of the time (in seconds) it takes for an individual animal to fall off the rotating rod, as well as the speed (in RPM) at which it falls and the distance travelled (in centimetres) during the test session. Before recording final observations, mice in all the groups were trained for three consecutive days. Each mouse was allowed a maximum of 180 seconds on the apparatus during the test recordings and acceleration of the rotating rod was gradually increased to 20 RPM over a course of 180 seconds (Graham and Sidhu, 2010).

### Wire hanging test

This test is used to analyse muscular strength of a mouse in terms of its ability to fall off (in seconds) a metal wire. Mice in all the groups were trained for three consecutive days before the initiation of the experiment. The total number of falls and reachings were recorded over a period of 180 seconds in all the groups (Ekmark-Lewén et al., 2018).

### Statistical analysis

All values are expressed as mean ± SEM of 10 animals in each experimental group. Data was statistically analyzed for p-values following one way analysis of variance and NK-test for comparisons between the experimental groups using GraphPad Prism 8.0 statistical software (San Diego, California, USA). Values with p<0.05 were considered as statistically significant.

### Tissue processing and histological examination

On the last day after behavioral observations, mice (n=5) were transcardially perfused with ice-cold saline, followed by perfusion with phosphate buffered paraformaldehyde solution (4%, v/v). Isolated brains were postfixed overnight at room temperature in 4% paraformaldehyde solution. Later, mid-brain sections were processed for routine Haematoxylin and Eosin (H&E) staining to demonstrate the appearance of the neuronal morphology and cell nuclei under light microscopy (Kiernan, 1999).

## Notes

### Competing Interest Statement

The authors have declared no competing interest.

## References

Abeliovich, A., Schmitz, Y., Fariñas, I., Choi-Lundberg, D., Ho, W.-H., Castillo, P.E., Shinsky, N., Verdugo, J.M.G., Armanini, M., and Ryan, A. (2000). Mice lacking α-synuclein display functional deficits in the nigrostriatal dopamine system. Neuron 25, 239–252.

Alim, M.A., Ma, Q.-L., Takeda, K., Aizawa, T., Matsubara, M., Nakamura, M., Asada, A., Saito, T., xkaji, M., and Yoshii, M. (2004). Demonstration of a role for α-synuclein as a functional microtubule-associated protein. Journal of Alzheimer’s Disease 6, 435–442.

Bucciantini, M., Giannoni, E., Chiti, F., Baroni, F., Formigli, L., Zurdo, J., Taddei, N., Ramponi, G., Dobson, C.M., and Stefani, M. (2002). Inherent toxicity of aggregates implies a common mechanism for protein misfolding diseases. nature 416, 507–511.

Cascella, R., Chen, S.W., Bigi, A., Camino, J.D., Xu, C.K., Dobson, C.M., Chiti, F., Cremades, N., and Cecchi, C. (2021). The release of toxic oligomers from α-synuclein fibrils induces dysfunction in neuronal cells. Nature communications 12, 1–16.

Chamachi, N.G., and Chakrabarty, S. (2017). Temperature-induced misfolding in prion protein: evidence of multiple partially disordered states stabilized by non-native hydrogen bonds. Biochemistry 56, 833–844.

Chandra, S., Gallardo, G., Fernández-Chacón, R., Schlüter, O.M., and Südhof, T.C. (2005). α- Synuclein cooperates with CSPα in preventing neurodegeneration. Cell 123, 383–396.

Chaudhary, K., Kumar, R., Singh, S., Tuknait, A., Gautam, A., Mathur, D., Anand, P., Varshney, G.C., and Raghava, G.P. (2016). A Web Server and Mobile App for Computing Hemolytic Potency of Peptides. Sci Rep 6, 22843.

Chu, D., Xu, W., Pan, R., Ding, Y., Sui, W., and Chen, P. (2015). Rational modification of oligoarginine for highly efficient siRNA delivery: structure–activity relationship and mechanism of intracellular trafficking of siRNA. Nanomedicine: Nanotechnology, Biology and Medicine 11, 435–446.

Clark, R.J., Fischer, H., Dempster, L., Daly, N.L., Rosengren, K.J., Nevin, S.T., Meunier, F.A., Adams, D.J., and Craik, D.J. (2005). Engineering stable peptide toxins by means of backbone cyclization: stabilization of the α-conotoxin MII. Proceedings of the National Academy of Sciences 102, 13767–13772.

Dawson, T.M., and Dawson, V.L. (2003). Rare genetic mutations shed light on the pathogenesis of Parkinson disease. The Journal of clinical investigation 111, 145–151.

Derossi, D., Chassaing, G., and Prochiantz, A. (1998). Trojan peptides: the penetratin system for intracellular delivery. Trends in cell biology 8, 84–87.

Di Giovanni, S., Eleuteri, S., Paleologou, K.E., Yin, G., Zweckstetter, M., Carrupt, P.-A., and Lashuel, H.A. (2010). Entacapone and tolcapone, two catechol O-methyltransferase inhibitors, block fibril formation of α-synuclein and β-amyloid and protect against amyloid-induced toxicity. Journal of Biological Chemistry 285, 14941-14954.

Drin, G., Cottin, S., Blanc, E., Rees, A.R., and Temsamani, J. (2003). Studies on the internalization mechanism of cationic cell-penetrating peptides. Journal of Biological Chemistry 278, 31192–31201.

Ekmark-Lewén, S., Lindström, V., Gumucio, A., Ihse, E., Behere, A., Kahle, P.J., Nordström, E., Eriksson, M., Erlandsson, A., and Bergström, J. (2018). Early fine motor impairment and behavioral dysfunction in (Thy-1)-h [A30P] alpha-synuclein mice. Brain and behavior 8, e00915.

Elmquist, A., Lindgren, M., Bartfai, T., and Langel, Ü. (2001). VE-cadherin-derived cell-penetrating peptide, pVEC, with carrier functions. Experimental cell research 269, 237–244.

Feni, L., and Neundorf, I. (2017). The Current Role of Cell-Penetrating Peptides in Cancer Therapy. Adv Exp Med Biol 1030, 279–295.

Franco, R., Rivas-Santisteban, R., Navarro, G., Pinna, A., and Reyes-Resina, I. (2021). Genes implicated in familial Parkinson’s disease provide a dual picture of nigral dopaminergic neurodegeneration with mitochondria taking center stage. International journal of molecular sciences 22, 4643.

Frankel, A., and Pabo, C. (1988). Cellular uptake of the Tat protein from human immunodeficiency virus. Cell 55, 1189-l.

Gautam, A., Chaudhary, K., Kumar, R., and Raghava, G.P.S. (2015a). Computer-aided virtual screening and designing of cell-penetrating peptides. In Cell-penetrating peptides (Springer), pp. 59–69.

Gautam, A., Chaudhary, K., Kumar, R., Sharma, A., Kapoor, P., Tyagi, A., and Raghava, G.P. (2013). In silico approaches for designing highly effective cell penetrating peptides. J Transl Med 11, 74.

Gautam, A., Nanda, J.S., Samuel, J.S., Kumari, M., Priyanka, P., Bedi, G., Nath, S.K., Mittal, G., Khatri, N., and Raghava, G.P. (2016). Topical Delivery of Protein and Peptide Using Novel Cell Penetrating Peptide IMT-P8. Sci Rep 6, 26278.

Gautam, A., Sharma, M., Vir, P., Chaudhary, K., Kapoor, P., Kumar, R., Nath, S.K., and Raghava, G.P. (2015b). Identification and characterization of novel protein-derived arginine-rich cell-penetrating peptides. European Journal of Pharmaceutics and Biopharmaceutics 89, 93–106.

Graham, D.R., and Sidhu, A. (2010). Mice expressing the A53T mutant form of human alpha-synuclein exhibit hyperactivity and reduced anxiety-like behavior. Journal of neuroscience research 88, 1777–1783.

Gupta, S., Kapoor, P., Chaudhary, K., Gautam, A., Kumar, R., and Raghava, G.P. (2013). In silico approach for predicting toxicity of peptides and proteins. PLoS One 8, e73957.

Haque, R., Shamsuzzama, L.K., Sharma, T., Fatima, S., Jadiya, P., Siddiqi, M.I., and Nazir, A. (2020). Human insulin modulates α-synuclein aggregation via DAF-2/DAF-16 signalling pathway by antagonising DAF-2 receptor in C. elegans model of Parkinson’s disease. Oncotarget 11, 634.

Huang, H., Zhang, W., Liu, D., Liu, B., Chen, G., and Zhong, C. (2011). Effect of temperature on gas adsorption and separation in ZIF-8: A combined experimental and molecular simulation study. Chemical engineering science 66, 6297–6305.

Jadiya, P., Chatterjee, M., Sammi, S.R., Kaur, S., Palit, G., and Nazir, A. (2011). Sir-2.1 modulates ’calorie-restriction-mediated’ prevention of neurodegeneration in Caenorhabditis elegans: implications for Parkinson’s disease. Biochem Biophys Res Commun 413, 306–310.

Jin, H., and Clayton, D.F. (1997). Synelfin regulation during the critical period for song learning in normal and isolated juvenile zebra finches. Neurobiology of learning and memory 68, 271–284.

Johansson, H.J., El-Andaloussi, S., Holm, T., Mäe, M., Jänes, J., Maimets, T., and Langel, Ü. (2008). Characterization of a novel Cytotoxic cell-penetrating Peptide Derived from p14ARF protein. Molecular Therapy 16, 115–123.

Kang, Z., Ding, G., Meng, Z., and Meng, Q. (2019). The rational design of cell-penetrating peptides for application in delivery systems. Peptides 121, 170149.

Kiernan, J.A. (1999). Histological and histochemical methods: theory and practice. Shock 12, 479.

Krüger, R., Kuhn, W., Müller, T., Woitalla, D., Graeber, M., Kösel, S., Przuntek, H., Epplen, J.T., Schols, L., and Riess, O. (1998). AlaSOPro mutation in the gene encoding α-synuclein in Parkinson’s disease. Nature genetics 18, 106–108.

Kumar, L., Shamsuzzama, Jadiya, P., Haque, R., Shukla, S., and Nazir, A. (2018). Functional Characterization of Novel Circular RNA Molecule, circzip-2 and Its Synthesizing Gene zip-2 in C. elegans Model of Parkinson’s Disease. Mol Neurobiol 55, 6914–6926.

Kumar, V., Patiyal, S., Dhall, A., Sharma, N., and Raghava, G.P.S. (2021). B3Pred: A Random- Forest-Based Method for Predicting and Designing Blood-Brain Barrier Penetrating Peptides. Pharmaceutics 13.

Lashuel, H.A., Overk, C.R., Oueslati, A., and Masliah, E. (2013). The many faces of α- synuclein: from structure and toxicity to therapeutic target. Nature Reviews Neuroscience 14, 38–48.

Li, B., Ge, P., Murray, K.A., Sheth, P., Zhang, M., Nair, G., Sawaya, M.R., Shin, W.S., Boyer, D.R., and Ye, S. (2018). Cryo-EM of full-length α-synuclein reveals fibril polymorphs with a common structural kernel. Nature communications 9, 1–10.

Lin, Z., and van Gunsteren, W.F. (2013). Refinement of the application of the GROMOS 54A7 force field to β-peptides. Journal of Computational Chemistry 34, 2796–2805.

Mahul-Mellier, A.-L., Burtscher, J., Maharjan, N., Weerens, L., Croisier, M., Kuttler, F., Leleu, M., Knott, G.W., and Lashuel, H.A. (2020). The process of Lewy body formation, rather than simply α-synuclein fibrillization, is one of the major drivers of neurodegeneration. Proceedings of the National Academy of Sciences 117, 4971–4982.

Marley, J., Lu, M., and Bracken, C. (2001). A method for efficient isotopic labeling of recombinant proteins. J Biomol NMR 20, 71–75.

Milletti, F. (2012). Cell-penetrating peptides: classes, origin, and current landscape. Drug discovery today 17, 850–860.

Morris, M.C., Vidal, P., Chaloin, L., Heitz, F., and Divita, G. (1997). A new peptide vector for efficient delivery of oligonucleotides into mammalian cells. Nucleic acids research 25, 2730–2736.

Ngambenjawong, C., Pineda, J.M.B., and Pun, S.H. (2016). Engineering an affinity-enhanced peptide through optimization of cyclization chemistry. Bioconjugate chemistry 27, 2854–2862.

Ngo, K.H., Yang, R., Das, P., Nguyen, G.K., Lim, K.W., Tam, J.P., Wu, B., and Phan, A.T. (2020). Cyclization of a G4-specific peptide enhances its stability and G-quadruplex binding affinity. Chemical Communications 56, 1082–1084.

Polymeropoulos, M.H., Lavedan, C., Leroy, E., Ide, S.E., Dehejia, A., Dutra, A., Pike, B., Root, H., Rubenstein, J., and Boyer, R. (1997). Mutation in the α-synuclein gene identified in families with Parkinson’s disease. science 276, 2045–2047.

Prochiantz, A. (1999). Homeodomain-derived peptides. In and out of the cells. Ann N Y Acad Sci 886, 172–179.

Qin, Y., Chen, H., Zhang, Q., Wang, X., Yuan, W., Kuai, R., Tang, J., Zhang, L., Zhang, Z., and Zhang, Q. (2011). Liposome formulated with TAT-modified cholesterol for improving brain delivery and therapeutic efficacy on brain glioma in animals. International journal of pharmaceutics 420, 304–312.

Qin, Y., Zhang, Q., Chen, H., Yuan, W., Kuai, R., Xie, F., Zhang, L., Wang, X., Zhang, Z., and Liu, J. (2012). Comparison of four different peptides to enhance accumulation of liposomes into the brain. Journal of drug targeting 20, 235–245.

Reissmann, S. (2014). Cell penetration: scope and limitations by the application of cell-penetrating peptides. Journal of Peptide Science 20, 760–784.

Scheller, A., Wiesner, B., Melzig, M., Bienert, M., and Oehlke, J. (2000). Evidence for an amphipathicity independent cellular uptake of amphipathic cell-penetrating peptides. Eur J Biochem 267, 6043–6050.

Schmidt, G.A., Jungclaus, J.H., Ammann, C., Bard, E., Braconnot, P., Crowley, T., Delaygue, G., Joos, F., Krivova, N., and Muscheler, R. (2011). Climate forcing reconstructions for use in PMIP simulations of the last millennium (v1. 0). Geoscientific Model Development 4, 33–45.

Shin, M.C., Zhang, J., Min, K.A., Lee, K., Byun, Y., David, A.E., He, H., and Yang, V.C. (2014). Cell-penetrating peptides: achievements and challenges in application for cancer treatment. J Biomed Mater Res A 102, 575–587.

Skwarczynski, M., and Toth, I. (2019). Cell-penetrating peptides in vaccine delivery: facts, challenges and perspectives (Future Science), pp. 465–467.

Soomets, U., Lindgren, M., Gallet, X., Hällbrink, M., Elmquist, A., Balaspiri, L., Zorko, M., Pooga, M., Brasseur, R., and Langel, U. (2000). Deletion analogues of transportan. Biochim Biophys Acta 1467, 165–176.

Stalmans, S., Bracke, N., Wynendaele, E., Gevaert, B., Peremans, K., Burvenich, C., Polis, I., and De Spiegeleer, B. (2015). Cell-penetrating peptides selectively cross the blood-brain barrier in vivo. PloS one 10, e0139652.

Stiernagle, T. (1999). Maintenance of C. elegans.

Tuttle, M.D., Comellas, G., Nieuwkoop, A.J., Covell, D.J., Berthold, D.A., Kloepper, K.D., Courtney, J.M., Kim, J.K., Barclay, A.M., and Kendall, A. (2016a). Solid-state NMR structure of a pathogenic fibril of full-length human α-synuclein. Nature structural & molecular biology 23, 409–415.

Tuttle, M.D., Comellas, G., Nieuwkoop, A.J., Covell, D.J., Berthold, D.A., Kloepper, K.D., Courtney, J.M., Kim, J.K., Barclay, A.M., Kendall, A., et al. (2016b). Solid-state NMR structure of a pathogenic fibril of full-length human α-synuclein. Nat Struct Mol Biol 23, 409–415.

van Raaij, M.E., Segers-Nolten, I.M., and Subramaniam, V. (2006). Quantitative morphological analysis reveals ultrastructural diversity of amyloid fibrils from α-synuclein mutants. Biophysical journal 91, L96–L98.

Vivès, E., Brodin, P., and Lebleu, B. (1997). A truncated HIV-1 Tat protein basic domain rapidly translocates through the plasma membrane and accumulates in the cell nucleus. J Biol Chem 272, 16010–16017.

Wender, P.A., Mitchell, D.J., Pattabiraman, K., Pelkey, E.T., Steinman, L., and Rothbard, J.B. (2000). The design, synthesis, and evaluation of molecules that enable or enhance cellular uptake: peptoid molecular transporters. Proc Natl Acad Sci U S A 97, 13003–13008.

Zarranz, J.J., Alegre, J., Gómez-Esteban, J.C., Lezcano, E., Ros, R., Ampuero, I., Vidal, L., Hoenicka, J., Rodriguez, O., and Atarés, B. (2004). The new mutation, E46K, of α-synuclein causes parkinson and Lewy body dementia. Annals of Neurology: Official Journal of the American Neurological Association and the Child Neurology Society 55, 164–173.

Zhang, Y., Guo, P., Ma, Z., Lu, P., Kebebe, D., and Liu, Z. (2021). Combination of cell-penetrating peptides with nanomaterials for the potential therapeutics of central nervous system disorders: a review. Journal of Nanobiotechnology 19, 1–22.

