## supplement file for "Penetratin inhibits α-synuclein fibrillation and improves locomotor functions in mice model of Parkinson’s disease"

### Supplementary Data

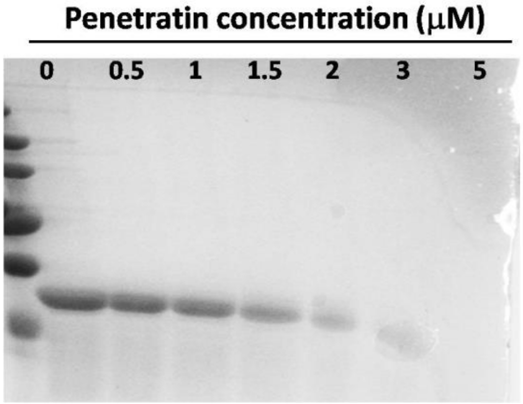


**Figure S1. Penetratin inhibits the α-Syn fibrillation in dose dependent manner. (A)** The α-syn fibrillation reaction was monitored until 18h. The reaction mixture was collected and ultra-centrifuged to separate supernatant and pellet. The pellet fraction was collected and loaded onto 15% SDS-PAGE. As seen, the α-syn amount decreases in pellet with increase in penetratin concentration.

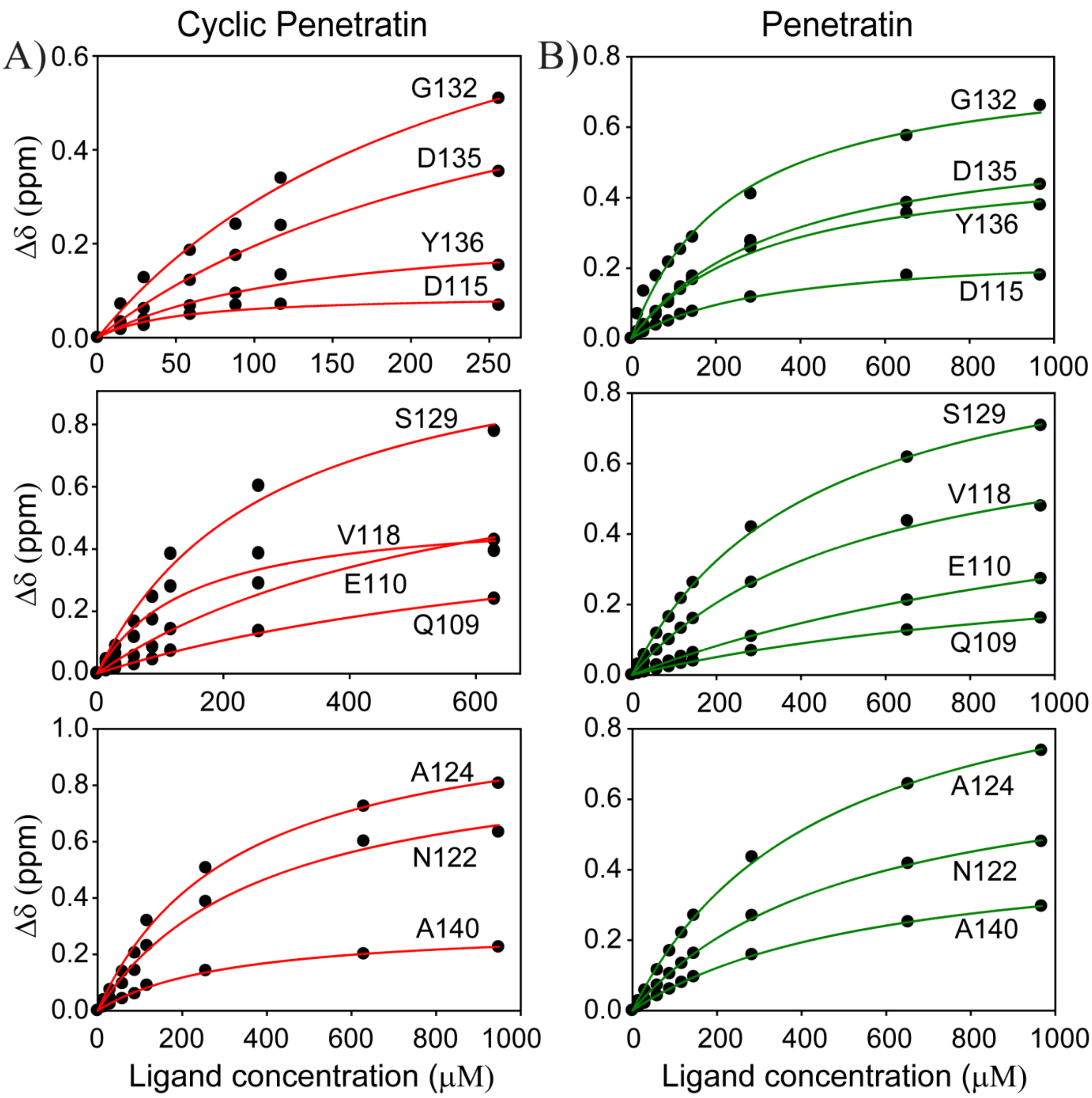


**Figure S2: Chemical shift perturbations (Δδ)** of C-terminus cross amide peaks showing perturbation obtained from $^{1}H$-$^{15}N$ HSQC spectra of α-Syn in presence of varying peptide concentrations **v/s** peptide concentration for **A)** Cyclic-penetratin (left panel) and **B)** Penetratin (right panel). The curve was fitted to obtain the dissociation constant ($K_D$) for each C-terminus residue.

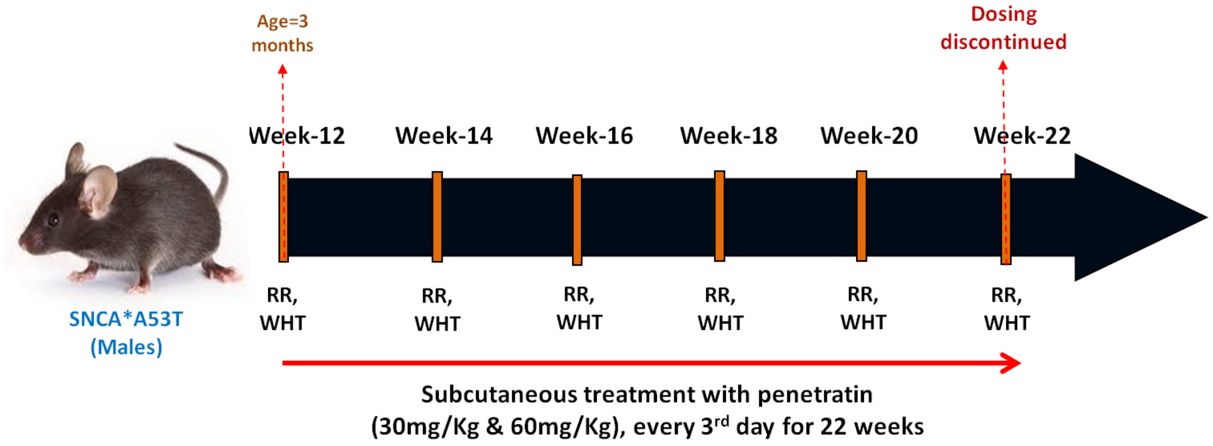


**Figure S3: Experimental plan for the mice behavioral studies.** PD mice were administered 30 or 60mg/kg of penetratin, every 3rd day beginning 12 weeks of age and continued until 22 weeks. The peptide was administered subcutaneously to PD mice and locomotor coordination activity was assessed using rotarod and wire hanging tests**.**

**Table S1:** $K_D$ value of C-terminus of α-Syn involved in the interaction with peptides

| Residues | Cyclic-Penetratin $K_D$ (μM) | Penetratin $K_D$ (μM) |
|---|---|---|
| Q109 | 899.13 ± 165.74 | 1261.57 ± 48.51 |
| E110 | 608.47 ± 178.05 | 1462.48 ± 47.69 |
| G111 | 101.88 ± 28.78 | 236.05 ± 35.18 |
| I112 | 230.11 ± 110.63 | 339.15 ± 27.77 |
| L113 | - | 466.56 ± 75.16 |
| E114 | 105.51 ± 37.53 | - |
| D115 | 51.22 ± 37.53 | 326.85 ± 33.68 |
| M116 | 566.10 ± 126.76 | 746.86 ± 43.92 |
| V118 | 146.25 ± 48.71 | 558.18 ± 45.48 |
| D119 | 214.47 ± 43.35 | 587.02 ± 19.71 |
| D121 | 311.72 ± 47.37 | 548.89 ± 27.49 |
| N122 | 395.47 ± 47.37 | 524.63 ± 23.85 |
| E123 | - | 658.07 ± 116.73 |
| A124 | 327.66 ± 37.77 | 451.57 ±23.39 |
| Y125 | 593.33 ± 176.84 | 422.45 ± 21.88 |
| E126 | 172.23 ± 67.25 | 488.30 ± 21.94 |
| M127 | 187.18 ± 76.72 | 286.02 ± 44.54 |
| S129 | 272.98 ± 54.78 | 435.04 ± 18.53 |
| E130 | 681.85 ± 323.27 | 423.03 ± 19.42 |
| E131 | 254.67 ± 237.43 | 678.42 ± 122.85 |
| G132 | 240.87 ± 51.30 | 241.32 ± 29.18 |
| Y133 | 188.60 ± 96.39 | 410.40 ± 34.44 |
| Q134 | 117.73 ± 32.18 | 510.92 ± 18.42 |
| D135 | 293.19 ± 61.33 | 365.93 ± 22.47 |
| Y136 | 135.33 ± 45.57 | 320.93 ± 31.06 |
| E137 | 71.51 ± 27.58 | 490.10 ± 27.70 |
| E139 | 258.62 ± 60.16 | 443.17 ± 26.26 |
| A140 | 312.20 ± 28.30 | 583.05 ± 16.90 |

**Table S2:** Effect of penetratin administration on motor coordination in PD transgenic animals assessed using Rotarod apparatus

| | Fall off time (in seconds) | | | | Acceleration (as RPM) | | | | Distance travelled (in centimeters) | | | |
|---|---|---|---|---|---|---|---|---|---|---|---|---|
| **Weeks** | **WT** | **PC** | **30mg/Kg** | **60mg/Kg** | **WT** | **PC** | **30mg/Kg** | **60mg/Kg** | **WT** | **PC** | **30mg/Kg** | **60mg/Kg** |
| **12** | 48.5±4.6 | 25.4±6.4[a] | 20.5±5.2 | 33.4±5.0 | 8.2±0.7 | 4.5±1.0[a] | 3.5±0.8 | 5.6±0.8 | 36.0±6.1 | 14.6±7.8[a] | 9.7±3.8 | 19.4±5.1 |
| **14** | 64.3±3.7 | 12.9±1.0[a] | 22.2±3.1[b] | 30.4±2.6[c] | 10.8±0.6 | 2.1±0.2[a] | 3.9±0.5[b] | 5.2±0.4[c] | 58.7±6. | 2.5±0.3[a] | 8.7±2.2 | 14.8±2.3[c] |
| **16** | 72.4±4.1 | 13.4±1.0[a] | 19.3±1.9 | 37.6±6.2[c] | 9.8±0.5 | 1.8±0.1[a] | 2.6±0.2 | 5.6±0.6[c] | 58.5±5.5 | 2.5±0.2[a] | 4.5±0.8 | 18.7±4.7[c] |
| **18** | 58.1±5.3 | 11.1±1.3[a] | 16.1±1.3 | 35.5±7.9[c] | 7.4±0.6 | 1.5±0.1[a] | 2.1±0.1 | 4.5±1.0[c] | 38.1±6.5 | 1.5±0.2[a] | 3.4±0.6 | 19.2±7.7[c] |
| **20** | 64.7±8.3 | 7.4±1.5[a] | 15.2±1.1 | 44.1±7.4[c] | 8.2±1.0 | 1.0±0.2[a] | 2.0±0.1 | 5.6±0.9[c] | 50.1±12.6 | 0.9±0.2[a] | 2.8±0.5 | 25.9±7.9[c] |
| **22** | 67.4±6.8 | 6.9±1.0[a] | 15.6±1.3 | 39.9±4.4[c] | 8.5±0.8 | 0.9±0.1[a] | 2.0±0.1 | 5.0±0.5[c] | 51.9±8.2 | 0.4±0.1[a] | 2.8±0.5 | 18.5±4.3[c] |

The values are expressed as mean±SEM, n=10. P<0.05, [a]significantly different from WT group,[b, c] significantly different from PC group.

**Table S3**: Effect of penetratin administration on locomotor function in PD transgenic animals assessed using wire-hanging test

| Reachings | | | | | Falls | | | |
|---|---|---|---|---|---|---|---|---|
| **Weeks** | **WT** | **PC** | **30mg/Kg** | **60mg/Kg** | **WT** | **PC** | **30mg/Kg** | **60mg/Kg** |
| **12** | 10.0±0.4 | 3.1±0.4[a] | 2.0±0.2[b] | 1.9±0.3[c] | 0 | 10.1±0.5[a] | 9.1±0.5 | 10.1±0.8 |
| **14** | 13.0±0.6 | 1.9±0.5[a] | 3.4±0.1[b] | 4.1±0.4[c] | 0 | 9.9±0.6[a] | 7.6±0.5[b] | 4.8±0.53[c] |
| **16** | 11.8±0.5 | 1.7±0.2[a] | 2.8±0.2 | 5.1±0.5[c] | 0 | 10.1±0.4[a] | 7.4±0.4[b] | 4.2±0.3[c] |
| **18** | 10.9±0.6 | 1.6±0.2[a] | 2.4±0.1 | 5.0±0.6[c] | 0 | 10.7±0.6[a] | 7.5±0.2[b] | 3.6±0.4[c] |
| **20** | 10.3±0.4 | 1.2±0.3[a] | 2.5±0.2 | 4.4±0.6[c] | 0 | 9.9±1.2[a] | 7.1±0.3[b] | 3.9±0.8[c] |
| **22** | 10.3±0.5 | 1.2±0.5[a] | 2.9±0.4 | 4.5±0.4[c] | 0 | 11.0±0.6[a] | 7.0±0.3[b] | 3.7±0.5[c] |

Values are expressed as mean±SEM, n=10. P<0.05, [a]significantly different from WT group,[b, c] significantly different from PC group.

**Table S4:** Potential number of H-bonds and salt-bridges between α-syn and peptides (n=8), analyzed from protein-peptides complex extracted at t=80 ns.

| **Protein** | **Peptide #** | **H-bonds** | **Salt-bridges** |
|---|---|---|---|
| **α-syn** | 1 | 3 | 1 |
| | 2 | 1 | 1 |
| | 3 | 0 | 0 |
| | 4 | 0 | 2 |
| | 5 | 2 | 0 |
| | 6 | 5 | 2 |
| | 7 | 1 | 0 |
| | 8 | 3 | 1 |

**Table S5.** Residues involved in H-bond (Black) and Salt-bridges (Blue) between α-syn and peptides.

| α-syn | Distance (nm) | Peptides |
|---|---|---|
| B:ASN 122[ ND2] | 2.83 | H:ALA 2[ O ] |
| B:LYS 80[ H ] | 1.71 | H:HIS 5[ O ] |
| B:ASN 122[ OD1] | 1.78 | H:ALA 2[ H ] |
| B:GLU 83[ OE1] | 3.96 | H:HIS 6[ NE2] |
| B:PRO 117[ O ] | 3.36 | G:SER 1[ OG ] |
| B:PRO 117[ O ] | 1.70 | G:ALA 2[ H ] |
| B:GLU 105[ OE1] | 1.99 | G:THR 4[ H ] |
| B:GLU 105[ OE2] | 1.86 | G:HIS 5[ H ] |
| B:GLU 105[ OE2] | 2.02 | G:HIS 6[ H ] |
| B:ASP 119[ OD1] | 3.40 | G:SER 1[ N ] |
| B:GLU 105[ OE2] | 3.99 | G:HIS 5[ NE2] |
| B:ASP 135[ H ] | 2.23 | C:SER 1[ O ] |
| B:TYR 133[ O ] | 1.93 | C:CYS 3[ H ] |
| B:ASP 115[ OD2] | 1.86 | K:ALA 2[ H ] |
| B:ASP 115[ OD1] | 3.69 | K:SER 1[ N ] |
| B:GLU 35[ OE1] | 3.01 | I:HIS 8[ NE2] |
| B:GLU 35[ OE2] | 2.97 | I:HIS 8[ NE2] |
